# DDIAS is a single-stranded DNA-binding effector of the TOPBP1-CIP2A complex in mitosis

**DOI:** 10.1101/2025.09.09.675193

**Authors:** Kaima Tsukada, Liudmyla Lototska, Aya Tsukada, Ipek Ilgin Gönenç, Charlotte Sherlaw-Sturrock, Manil Kanade, Antony W. Oliver, Samuel E. Jones, Satpal S. Jhujh, Julius Bannister, Asmat Ali, Muhammad Raza, Mathias Toft, Zafar Iqbal, Ambrin Fatima, Thomas C. R. Miller, Grant S. Stewart, Fena Ochs, Andrew N. Blackford

## Abstract

DNA double-strand breaks and unresolved DNA replication intermediates are particularly dangerous during mitosis. Paradoxically, cells inactivate canonical DNA repair mechanisms during chromosome segregation in favor of alternative pathways that depend on TOPBP1 and CIP2A, but how they function is still poorly defined. Here, we describe the identification of DDIAS as a mitosis-specific DNA damage response protein. We establish DDIAS as a phosphorylation-dependent component and effector of the TOPBP1-CIP2A complex with single-stranded DNA (ssDNA)-binding activity, and thereby delineate a ssDNA protection mechanism that safeguards chromosome integrity during mitosis, particularly in BRCA-deficient cells. We also identify biallelic inactivating mutations in *DDIAS* in a patient with a severe neurodevelopmental disorder. These findings highlight for the first time a potential physiological role for the DNA damage response in mitosis.

## INTRODUCTION

Maintenance of genome stability is fundamental for life, as DNA damage and chromosomal aberrations can lead to premature senescence, cell death and disease.^1^ This is highlighted by individuals with rare inherited chromosome instability disorders such as Bloom syndrome, Fanconi anaemia, and Nijmegen breakage syndrome, which are characterized by phenotypes including microcephaly, immunodeficiency, and increased cancer predisposition.^2^ Cells have therefore evolved a robust DNA damage response network to detect and orchestrate repair of DNA lesions.^3^ During interphase, DNA damage signalling can trigger checkpoint activation and cell cycle arrest, and ultimately senescence or apoptosis if lesions cannot be repaired.^1^ This ensures that cells do not enter mitosis with broken chromosomes, which would be catastrophic for genome stability and cell survival.^4^

Mitosis presents a unique challenge for genome stability maintenance, because once cells have committed to division, DNA damage checkpoints and most DNA repair pathways are inactivated until the following G1 phase.^4^ In addition, mitotic chromatin becomes highly condensed, rendering DNA lesions less accessible and more difficult to repair; nonetheless, it has become clear that cells do mount a mitosis-specific DNA damage response, which is crucial for preventing chromosome missegregation and micronucleus formation. The initial stages of this response resemble those in interphase,^1^ where DNA double-strand breaks (DSBs) activate the ATM kinase, which phosphorylates histone H2AX and recruits the mediator protein MDC1. This leads to recruitment of the tumour suppressors BRCA1 and 53BP1 in interphase cells, followed by rapid and accurate DNA repair via homologous recombination or non-homologous end-joining. But in mitosis, BRCA1 and 53BP1 recruitment is blocked,^5–8^ and instead a complex containing TOPBP1 and CIP2A is recruited to DNA lesions by MDC1.^9–12^ These TOPBP1-CIP2A complexes form filamentous assemblies that can bridge discrete MDC1 foci.^9^ This has led to the hypothesis that instead of repairing breaks, TOPBP1 and CIP2A tether broken DNA ends during mitosis to ensure that the two chromosome fragments segregate into one daughter nucleus, where they can be repaired accurately and efficiently after chromosome decondensation.^4^ Two elegant studies subsequently provided clear evidence for this, demonstrating that TOPBP1 and CIP2A promote reassembly of shattered chromosomes in a model of chromothripsis in cancer cells.^13,14^

More recently, it has been revealed that error-prone DNA repair pathways including microhomology-mediated end-joining (MMEJ) are active in mitosis, suggesting that limited DNA repair can in fact take place during this cell cycle phase.^15–20^ In addition, cells have mechanisms in mitosis to unwind or cleave sister chromatid entanglements caused by incomplete DNA replication during the previous S phase, which would otherwise prevent sister chromatid disjunction.^4,21^ Recent evidence indicates that TOPBP1 and CIP2A may play a role in some or all of these processes, although this is still unclear due to conflicting observations.^19,20,22–30^

Understanding how cells protect damaged or incompletely replicated chromosomes during mitosis has important implications for human health, given that some cancers (especially those deficient in the breast/ovarian tumour suppressors BRCA1 and BRCA2) rely on it for survival.^10,11^ However, many key questions remain. Neither TOPBP1 nor CIP2A contain DNA-binding domains, suggesting that at least one additional factor might be required to link them to DNA lesions. We also have limited understanding as to how the mitotic DNA damage response is regulated. Moreover, despite the identification of numerous human syndromes associated with defects in interphase DNA repair pathways,^2^ to date no inherited diseases have been linked to aberrant mitotic DNA damage responses. Consequently, the pathological impact of disrupting the TOPBP1-CIP2A pathway on human physiology is still unknown.

Here, we describe the identification of bi-allelic inactivating mutations in the *DDIAS* gene in a patient presenting with microcephaly, global developmental delay, seizures, intellectual disability, and other clinical features similar to those seen in DNA repair deficiency syndromes. We show that *DDIAS* encodes a previously unrecognised member of the mitotic DNA damage response with single-stranded DNA (ssDNA)-binding activity. CDK1 and PLK1 phosphorylate DDIAS to promote a high-affinity interaction with TOPBP1 during mitosis. Like CIP2A, ablation of DDIAS in human cells causes chromosomal instability and is synthetically lethal with BRCA1/2 deficiency. Our findings position DDIAS as the third core component of the mitotic TOPBP1-CIP2A network and delineate a phosphorylation-dependent DNA-end bridging mechanism that safeguards chromosome integrity during cell division.

## RESULTS

### *DDIAS* is mutated in a neurodevelopmental disorder with symptoms similar to those found in genome instability syndromes

We identified a five-year-old male child of South Asian heritage from a consanguineous marriage (**Figure 1A**), who started having frequent generalized clonic seizures one week after birth. The proband presented with severe neurodevelopmental and other abnormalities including microcephaly, short stature, global developmental delay, intellectual disability, limited speech and generalized spasticity (**Figures 1B and 1C**). Clinical observations are summarized in **Table 1** and described in detail in the Methods. Whole exome sequencing was carried out to determine the underlying cause of disease. This analysis identified a likely pathogenic homozygous nonsense variant in *DDIAS* (NM_145018.4:c.1631C>A; p.Ser544Ter), a poorly characterized gene that has been linked to control of apoptosis and co-evolved with homologous recombination genes (**Figure 1D**).^31,32^ Based on the phenotypic overlap of the proband’s clinical symptoms with those of chromosomal instability disorders,^2^ we wished to explore whether DDIAS plays a role in DNA repair. In line with this possibility, patient-derived cells displayed chromosomal instability in the form of increased micronucleation (**Figure 1E**) and chromosomal aberrations (**Figure 1F**). Importantly, the number of micronuclei was restored to levels observed in a healthy donor’s cells upon exogenous expression of DDIAS (**Figures 1G and S1A**), indicating that the chromosomal instability in the patient-derived cells is caused by lack of wild-type (WT) DDIAS expression.

**Figure 1.**
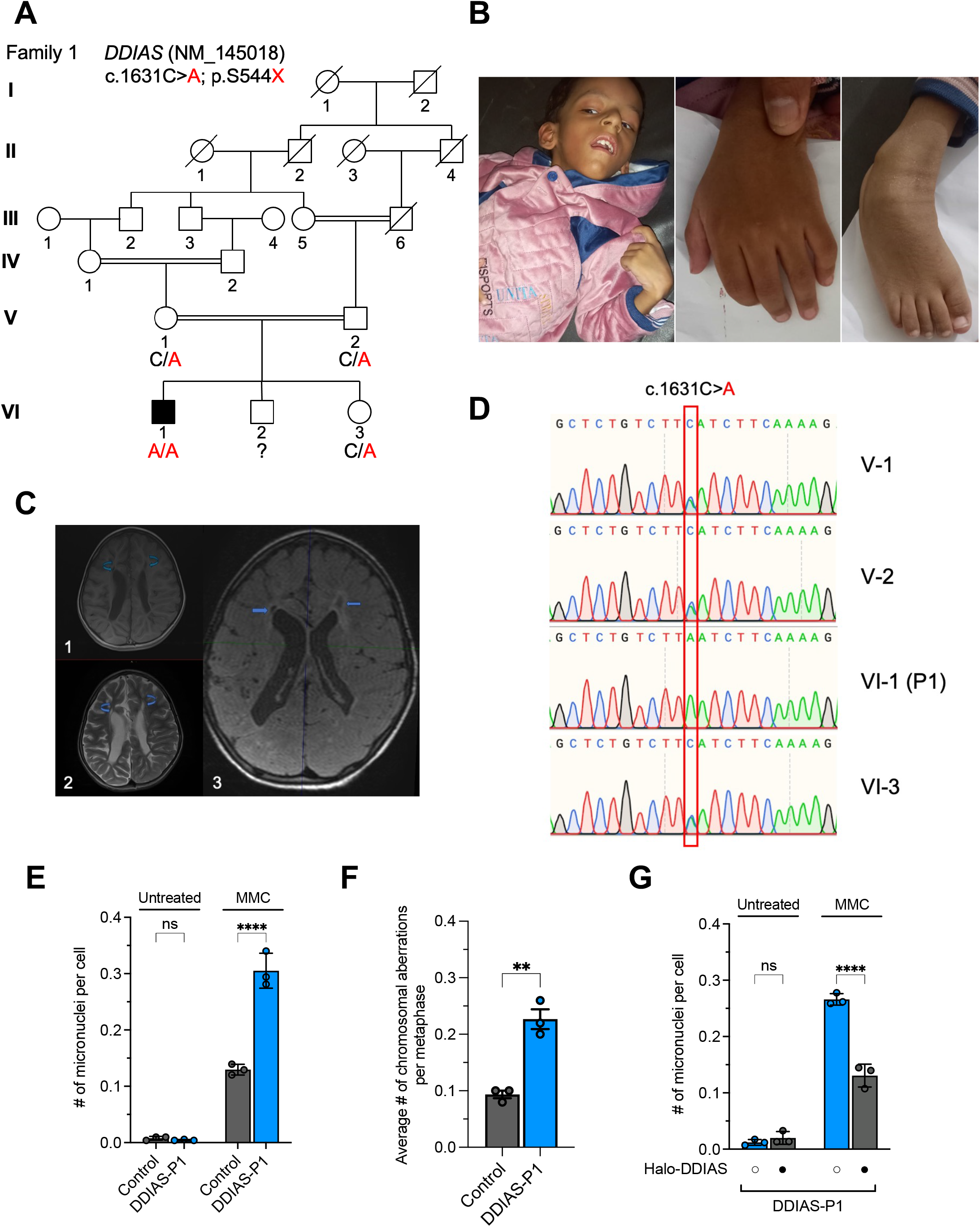
*DDIAS* is mutated in a patient with a neurodevelopmental disorder and chromosomal instability. **(A)** Pedigree of patient DDIAS-P1 in a family with a *DDIAS* inactivating variant. Filled square indicates the patient, and double horizontal line indicates consanguinity. Question mark denotes individual VI-2 has not undergone WES. **(B)** Photographs of patient DDIAS-P1. **(C)** MRI of patient’s brain, showing abnormal signals in white matter along bilateral frontal horns depicted by the arrows, appearing hypointense on T1 (1) and hyperintense on T2 weighted images (2). Abnormal signals in white matter along bilateral frontal horns depicted by the arrows, appearing hypointense on FLAIR images (3). **(D)** Chromatograms showing the segregation status of the *DDIAS* variant. **(E)** Quantification of micronuclei in patient-derived fibroblasts and cells from a healthy donor control. Where indicated, cells were treated with 50 ng/mL MMC for 48 hr. Error bars denote SD from n = 3 experiments. Un-paired two-tailed t-test was used to calculate significance. **(F)** Quantification of chromosomal aberrations in patient-derived fibroblasts and cells from a healthy donor control. Error bars denote SD from n = 3 experiments. Un-paired two-tailed t-test was used to calculate significance. **(G)** Quantification of micronuclei in patient-derived fibroblasts treated with 50 ng/mL MMC for 48 hr, complemented where indicated with exogenous DDIAS. Error bars denote SD from n = 3 experiments. Un-paired two-tailed t-test was used to calculate significance. See also **Figure S1**.

**Table 1.** Summary of clinical phenotypes.

|  | **DDIAS-P1** |
| --- | --- |
| **Gene** | *DDIAS* |
| **Allele 1** | NM_145018.4:c.1631C>A;<br>p.Ser544Ter |
| **Allele 2** | NM_145018.4:c.1631C>A;<br>p.Ser544Ter |
| **Origin** | South Asian |
| **Gender** | Male |
| **Age at last examination** | 5 years |
| **Patient phenotype** | Growth defects |
|  | Seizures |
|  | Microcephaly |
|  | Brain abnormalities on MRI |
|  | Global developmental delay |
|  | Intellectual disability |
|  | Limited speech |
|  | Facial dysmorphism |
|  | Generalized spasticity |
|  | Café-au-lait spots |
|  | Recurrent lung infections |
|  | Hypertonia |
|  | Horizontal nystagmus |
|  | High arched palate |
|  | Bilateral clinodactyly |

### DDIAS is a mitotic DNA damage response protein

Proteins involved in DNA repair processes often form foci at DNA lesions that can be observed by immunofluorescence microscopy to colocalize with phosphorylated H2AX (γH2AX).^33^ Endogenous DDIAS was not detectable using commercially available antibodies; we therefore generated *DDIAS* knockout RPE-1 and DLD-1 cells using CRISPR-Cas9, and established a doxycycline-inducible HaloTag-fused DDIAS expression system via lentiviral transduction to enable us to visualize DDIAS localization (**Figures S1B-S1E**). In interphase cells, DDIAS distribution was mostly pan-nuclear, and this did not change when cells were exposed to various genotoxic drugs (**Figures 2A and S1F**), including the DNA interstrand crosslinking agent mitomycin C (MMC), the radiomimetic drug neocarzinostatin (NCS), or the DNA polymerase inhibitor aphidicolin (APH). Strikingly however, we noticed that DDIAS accumulated in bright foci that colocalized with γH2AX in cells that were undergoing mitosis in the presence of genotoxic stress (**Figures 2A and S1F**).

**Figure 2.**
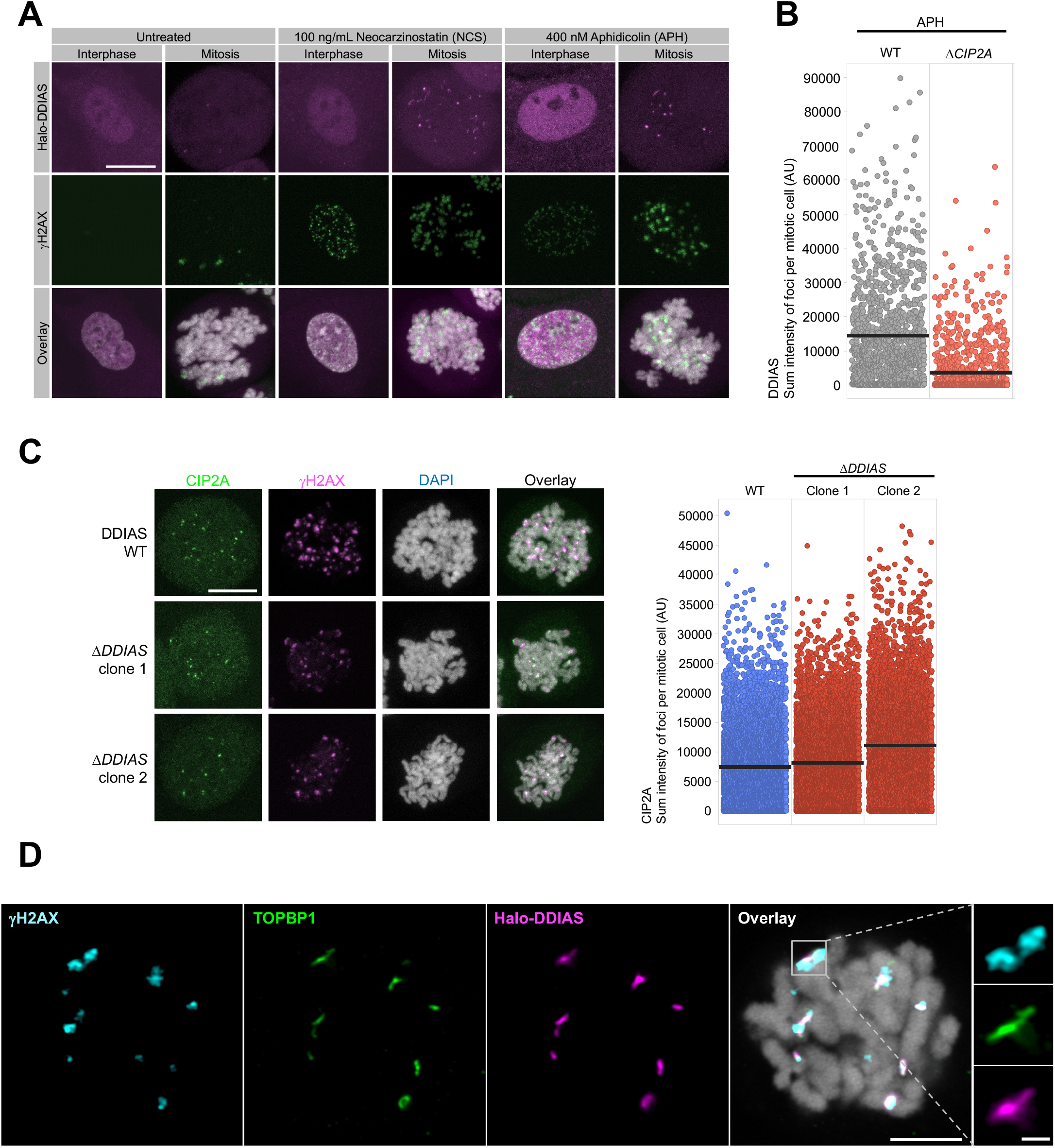
DDIAS is a mitotic DNA damage response protein. **(A)** Representative immunofluorescence images of DDIAS and γH2AX foci formation in Δ*DDIAS* RPE-1 cells expressing Halo-tagged DDIAS. Cells were treated with 100 ng/mL NCS for 30 min or 400 nM APH for 24 hr where indicated. DNA was stained with DAPI. Scale bar, 10 μm. **(B)** Quantification of the sum of DDIAS foci intensity per mitotic cell by QIBC in WT and Δ*CIP2A* RPE-1 cells treated with APH and nocodazole. n > 1200 cells for each cell line. Cells were gated by H3-pS10 intensity to quantify mitotic cells only. Means are shown as black bars. AU, arbitrary units. **(C)** Representative immunofluorescence images of CIP2A and γH2AX foci in two independent Δ*DDIAS* DLD-1 clones and parental DLD-1 (WT) cells treated with APH and nocodazole. DNA was stained with DAPI. Scale bar, 10 μm. Sum of CIP2A foci intensity per mitotic cell was quantified by QIBC. n > 7000 cells for each cell line. Cells were gated by H3-pS10 intensity to quantify mitotic cells only. Means are shown as black bars. **(D)** Airyscan super-resolution confocal single slice images of Halo-DDIAS, TOPBP1 and γH2AX foci in mitosis 1 hr after 0.5 Gy IR. DNA was stained with DAPI. Scale bar (main), 5 μm; scale bar (inset), 1 μm. See also **Figure S1**.

This observation indicated that DDIAS may play a role in response to DNA damage, but only during mitosis—a characteristic that is highly reminiscent of CIP2A.^10,12^ We therefore wished to determine whether DDIAS and CIP2A are involved in the same mitotic DNA damage response pathway. To do this, we first checked whether DDIAS recruitment was dependent on CIP2A, by examining Halo-DDIAS foci formation in WT and Δ*CIP2A* RPE-1 cells.^12^ Significantly, we observed that DDIAS failed to accumulate in foci following APH treatment in the absence of CIP2A (**Figure 2B**). Conversely, CIP2A and γH2AX foci formation were not defective in Δ*DDIAS* cells (**Figure 2C**), suggesting that DDIAS acts downstream of TOPBP1-CIP2A complex recruitment in the mitotic DNA damage response.

We have previously shown using high-resolution microscopy that TOPBP1 and CIP2A form filament-like structures at DNA lesions in mitosis, with TOPBP1 filaments being observed to bridge discrete ionizing radiation (IR)-induced foci occasionally—indicative of a DNA end-bridging role.^9,12^ We therefore assessed using the same technique where DDIAS localizes in relation to γH2AX and TOPBP1 at DSBs, and found that the DDIAS localization pattern closely resembles that of TOPBP1 rather than γH2AX, with DDIAS and TOPBP1 colocalizing at filamentous structures that sometimes bridge γH2AX foci (**Figure 2D**). These observations provide further support for a role for DDIAS in the TOPBP1-CIP2A pathway in mitosis.

### Mechanism of DDIAS recruitment to mitotic DNA lesions

DDIAS is a protein of 998 amino acid residues, which is predicted by AlphaFold 3^34^ to be largely unstructured except for an N-terminal oligonucleotide-binding (OB) fold (**Figure 3A**). The patient mutation S544X could potentially produce a truncated DDIAS protein containing the OB fold but lacking most of the C-terminal unstructured region. To test the requirement for these regions in mediating DDIAS recruitment to mitotic DNA lesions, we generated Δ*DDIAS* DLD-1 cells expressing either the S544X patient mutant or a DDIAS mutant lacking the OB fold (ΔOB; **Figure S1E**). Next, we examined the ability of these mutant proteins to form foci in mitotic cells, and found that while the ΔOB mutant behaved like the WT protein, the S544X mutant was profoundly defective in foci formation (**Figure 3B**). These results demonstrate that sequences in the C-terminal half of DDIAS direct its localization to DNA damage sites, but the OB fold is dispensable in this regard.

**Figure 3.**
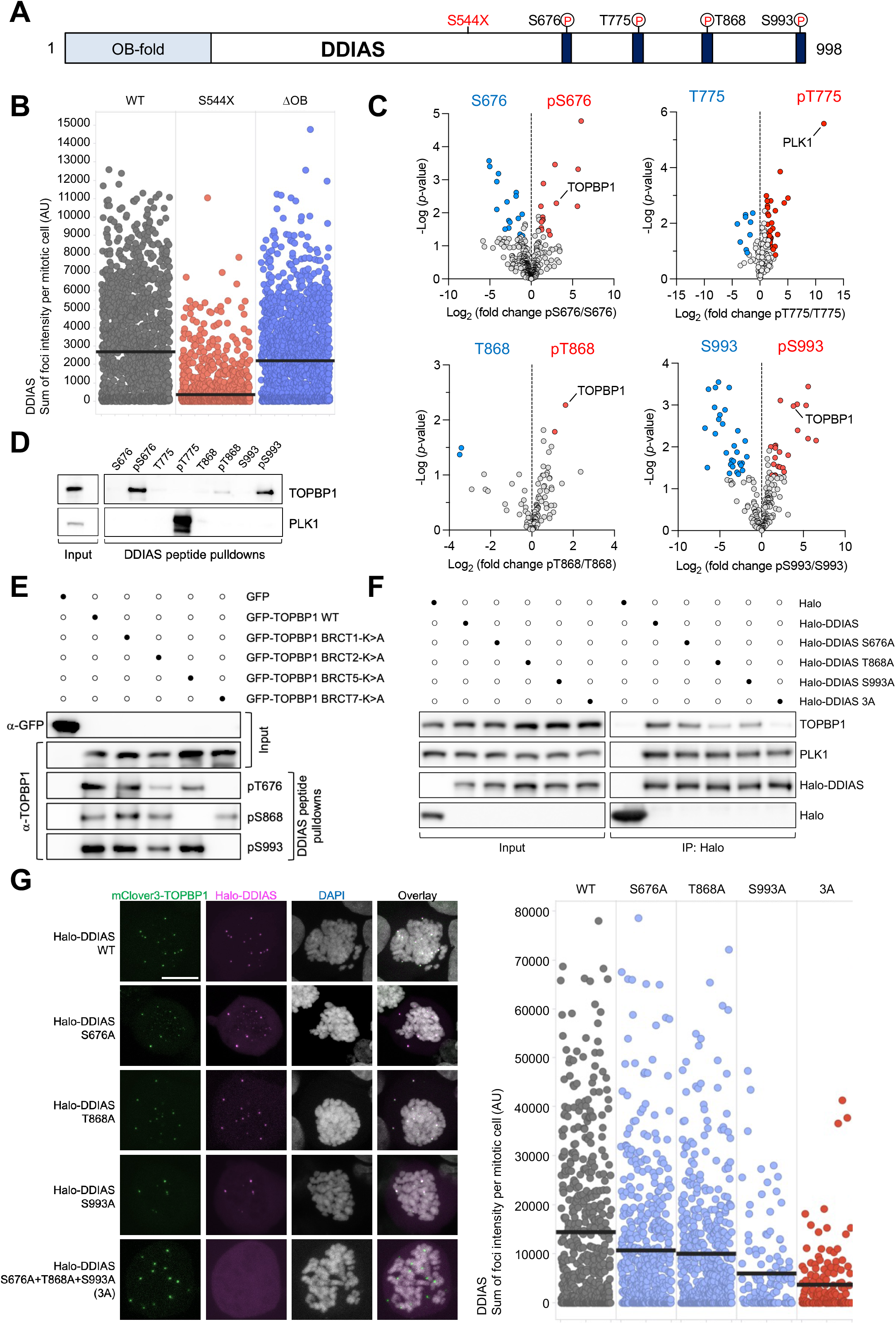
Mechanism of DDIAS recruitment to mitotic DNA lesions. **(A)** Schematic of the human DDIAS protein showing the N-terminal OB fold domain, the location of the patient mutation (S544X), and the conserved motifs in the unstructured C-terminus. **(B)** Quantification of the sum of DDIAS foci intensity per mitotic cell by QIBC in Δ*DDIAS* DLD-1 cells expressing the indicated Halo-tagged DDIAS proteins and treated with APH and nocodazole. n > 1200 cells for each cell line. Cells were gated by H3-pS10 intensity to quantify mitotic cells only. Means are shown as black bars. **(C)** Volcano plots depicting log_2_ fold enrichment and -log p-value of the indicated native versus phosphorylated DDIAS peptide-binding proteins from 3 independent experiments. See also **Tables S1-S4**. **(D)** Pulldowns from 293FT cell lysates using the indicated biotinylated native or phosphorylated DDIAS peptides. **(E)** Pulldowns from lysates of 293FT cells transfected with plasmids expressing GFP, GFP-tagged WT TOPBP1 or GFP-TOPBP1 BRCT domain mutants, using the indicated DDIAS phospho-peptides. **(F)** Immunoprecipitations from lysates of 293FT cells transiently transfected with plasmids expressing Halo or the indicated Halo-tagged DDIAS proteins. Cells were treated with 100 ng/mL nocodazole for 16 hr prior to harvesting. **(G)** Representative immunofluorescence images of U2OS cells treated with APH and nocodazole, and transiently transfected with plasmids expressing WT or the indicated mutant Halo-DDIAS proteins. DNA was stained with DAPI. Scale bar, 10 μm. Sum of DDIAS foci intensity per mitotic cell was quantified by QIBC. n > 300 cells for each condition. Cells were gated by H3-pS10 intensity to quantify mitotic cells only. Means are shown as black bars. See also **Figures S2-S5**.

Sequence alignment of vertebrate DDIAS orthologues revealed four highly conserved short linear motifs (SLiMs) within residues 544–998 (**Figures 3A and S2**), each containing potential Ser/Thr phosphorylation sites. Pulldowns from whole-cell extracts using biotinylated native and phosphorylated peptides derived from these SLiM sequences were therefore carried out to identify potential protein binding partners. Quantitative liquid chromatography-tandem mass spectrometry (LC-MS/MS) analyses showed that TOPBP1 was among the top hits in the pSer676, pThr868 and pSer993 peptide pulldowns, whereas the mitotic kinase PLK1 was the top hit in the pThr775 peptide pulldown (**Figure 3C**). We observed consistent results by western blotting (**Figure 3D**).

TOPBP1 contains 9 BRCT domains, 4 of which have phospho-peptide binding ability (**Figure S3A**).^35^ To determine whether one or more of these domains were responsible for binding DDIAS phospho-peptides, we expressed WT TOPBP1 and a set of BRCT mutant proteins in 293FT cells, each with one phosphate-binding lysine residue mutated to alanine to abolish phospho-peptide binding: BRCT1 (K154A), BRCT2 (K250A), BRCT5 (K704A) and BRCT7 (K1317A).^9,23^ Peptide pulldowns were carried out from lysates from these cells, which revealed that pSer676 and pSer993 peptide binding to TOPBP1 was abolished by mutation of BRCT domain 7, whereas pThr868 peptide binding was lost upon mutation of BRCT domain 5 (**Figure 3E**). In line with this, the primary amino acid sequences of the DDIAS Ser676 and Ser993 motifs are similar to the TOPBP1 BRCT7-binding motif in FANCJ (**Figure S3B**),^36^ and the DDIAS-Thr868 motif closely resembles the TOPBP1 BRCT5-binding motif in BLM (**Figure S3C**).^23,37^ Supporting results were provided by AlphaFold 3 structural predictions (**Figure S3D**), and fluorescence polarization studies (**Figures S4A and S4B**), which showed that the DDIAS pSer676 and pSer993 motifs selectively bind TOPBP1 BRCT domain 7 and the pThr868 motif binds BRCT domain 5, with dissociation constants in the nanomolar range.

Next, we examined the effect of mutating the 3 DDIAS phospho-sites alone or in combination in the context of the full-length protein. In initial experiments, we observed that DDIAS immunoprecipitates contain TOPBP1 and CIP2A, and that addition of nocodazole to arrest cells in early mitosis dramatically enhanced DDIAS-TOPBP1 interaction, in keeping with a role for DDIAS during cell division (**Figure S5A**). We then examined this interaction in pulldowns using Halo-DDIAS WT and phospho-site mutants S676A, T868A and S993A as baits. Interestingly, mutation of each of these sites individually had a mild but consistently detrimental effect on DDIAS-TOPBP1 interaction, whereas mutation of all three sites together in a “3A” mutant almost abrogated it (**Figure 3F**). Reciprocal pulldowns using GFP-TOPBP1 proteins as baits showed that BRCT5 and BRCT7 mutants reduced DDIAS binding, whereas a BRCT5/7 double mutant almost abolished the interaction (**Figure S5B**). We conclude based on these experiments that TOPBP1 and DDIAS interact directly in a manner dependent on multiple TOPBP1 BRCT domains and phospho-sites in DDIAS.

Finally, we investigated whether the TOPBP1-DDIAS interaction was required for DDIAS recruitment to mitotic DNA lesions. We therefore examined foci formation of the DDIAS S676A, T868A and S993A single mutants, as well as the 3A triple mutant. In line with our pulldown data (**Figure 3F**), we observed a partial recruitment defect with single mutants, whereas mutation of all three phospho-sites abolished DDIAS foci formation (**Figure 3G**). Thus, TOPBP1 recruits DDIAS to DNA lesions in mitosis via multivalent phospho-dependent interactions.

### CDK1 and PLK1 promote DDIAS binding to TOPBP1

LC-MS/MS analyses of the DDIAS-pThr775 peptide pulldown identified the mitotic kinase PLK1 as a top hit (**Figure 3C**). Notably, Thr775 and its surrounding residues match the consensus Ser-Thr-Pro binding sequence for the polo-box domain (PBD) of PLK1 (**Figure S2**).^38^ Support for a direct interaction between PLK1 and DDIAS was provided by AlphaFold 3, which predicted docking of the DDIAS-pThr775 motif with the PLK1-PBD (**Figure 4A**). To verify this interaction, we carried out pulldowns using Halo-DDIAS WT and a T775A mutant expressed in nocodazole-arrested 293FT cells. While we readily observed PLK1 binding to WT DDIAS, this interaction was barely detectable when Thr775 was mutated (**Figure 4B**). We conclude that when DDIAS is phosphorylated on Thr775 in cells, it provides a binding site for the PBD of PLK1.

**Figure 4.**
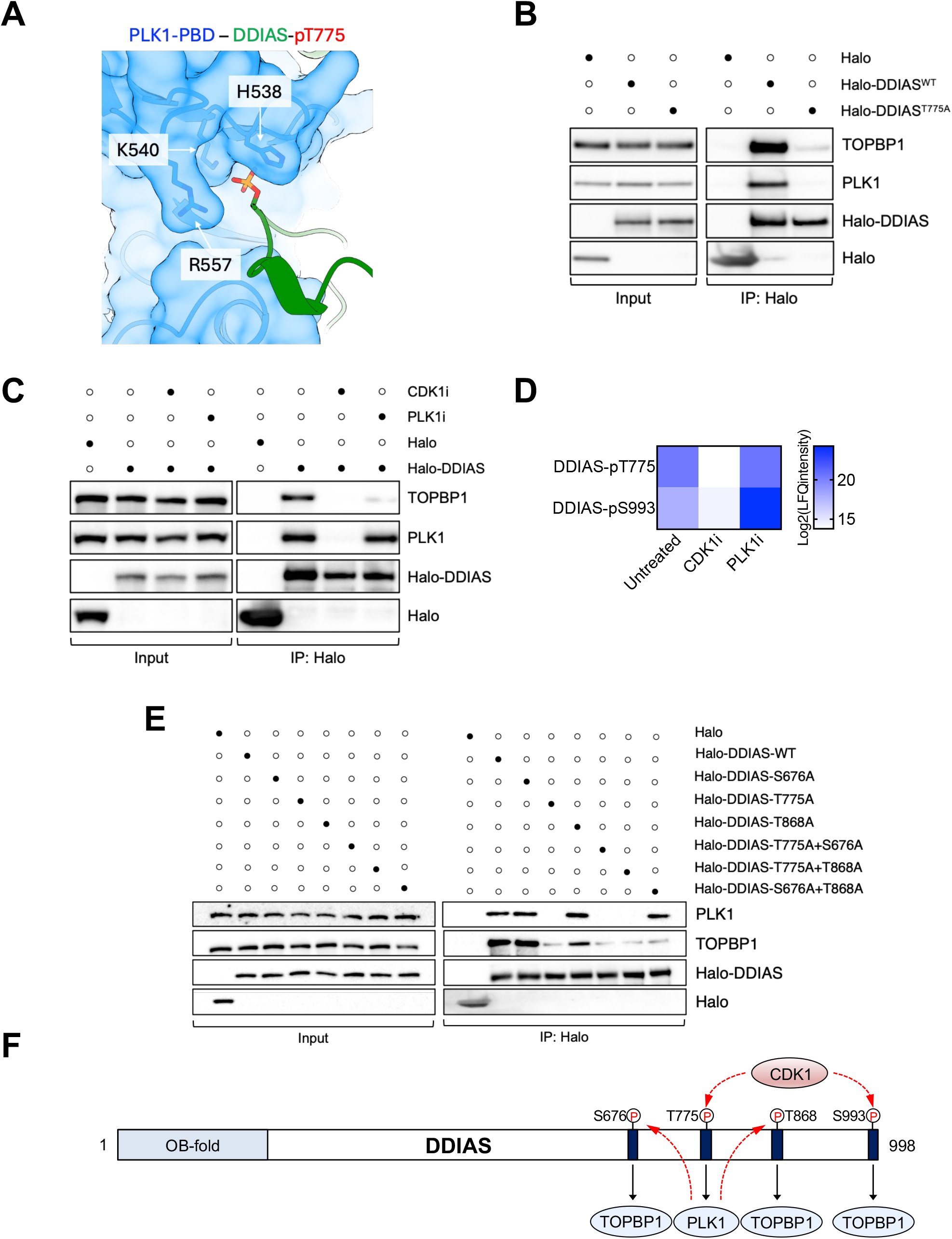
CDK1 and PLK1 promote DDIAS binding to TOPBP1. **(A)** AlphaFold 3 model of the direct interaction between DDIAS-pThr775 and the key phosphate-binding residues (His538, Lys540 and Arg557) in the PBD of PLK1. **(B)** Immunoprecipitations from lysates of 293FT cells transiently transfected with plasmids expressing Halo or the indicated Halo-tagged DDIAS proteins. Cells were treated with 100 ng/mL nocodazole for 16 hr prior to harvesting. **(C)** Immunoprecipitations from lysates of 293FT cells transiently transfected with plasmids expressing Halo or Halo-tagged DDIAS. Cells were treated with 100 ng/mL nocodazole for 16 hr prior to harvesting, and where indicated with 9 µM CDK1 inhibitor RO-3306 or 25 nM PLK1 inhibitor BI 2356 for 2 hr prior to harvesting. **(D)** Phospho-proteomics analyses of DDIAS phosphorylation on Thr775 and Ser993. Immunoprecipitations were carried out from lysates of 293FT cells transiently transfected with plasmids expressing Halo-DDIAS. Cells were treated with 100 ng/mL nocodazole for 16 hr prior to harvesting, and either 9 µM CDK1 inhibitor RO-3306 or 25 nM PLK1 inhibitor BI 2356 for 2 hr prior to harvesting as appropriate. See also **Table S5**. **(E)** Immunoprecipitations from lysates of 293FT cells transiently transfected with plasmids expressing Halo or the indicated Halo-tagged DDIAS proteins. Cells were treated with 100 ng/mL nocodazole for 16 hr prior to harvesting. **(F)** Model showing the conserved motifs in human DDIAS that are phosphorylated by CDK1 and PLK1, to promote DDIAS interactions with TOPBP1 and PLK1.

Interestingly, the DDIAS-T775A mutant was also defective in its interaction with TOPBP1 (**Figure 4B**), even though TOPBP1 was not detectable in the pThr775 peptide pulldown (**Figure 3D and Table S2**). This indicated that PLK1 might be responsible for phosphorylating one or more of the residues in DDIAS that bind to the TOPBP1 BRCT domains. CDK1 often acts as a priming kinase to generate substrates that are bound and subsequently phosphorylated by PLK1.^39^ We therefore tested how acute inhibition of CDK1 or PLK1 altered DDIAS complex assembly. CDK1 inhibition with RO-3306^40^ blocked interaction of DDIAS with both TOPBP1 and PLK1, whereas PLK1 inhibition with BI 2356^41^ led to decreased DDIAS-TOPBP1 binding but had little effect on the DDIAS-PLK1 interaction (**Figure 4C**). In line with this, phospho-proteomics of Halo-DDIAS isolated from nocodazole-treated cells revealed that CDK1 inhibition led to a reduction in Thr775 and Ser993 phosphorylation, whereas PLK1 inhibition did not (**Figure 4D**).

Unfortunately, we could not detect Ser676 or Thr868 phosphorylation in these analyses, so we could not assess the effect of CDK1 or PLK1 inhibition on modification of these sites. We therefore reasoned that if PLK1 recruitment via binding to DDIAS-pThr775 allows PLK1 to phosphorylate Ser676 and Thr868, then firstly a double S676A/T868A DDIAS mutant would phenocopy a T775A mutant in terms of deficiency in TOPBP1 binding; and secondly, T775A in combination with either S676A or T868A would be no more defective than a T775A mutation alone. This is indeed what we observed using single and double DDIAS mutants in further pulldown experiments (**Figure 4E**).

Taken together, these observations are consistent with a model in which CDK1 directly promotes DDIAS-TOPBP1 binding by phosphorylating DDIAS on Ser993; and indirectly by phosphorylating DDIAS on Thr775 to produce a binding site for PLK1. PLK1-DDIAS binding then promotes further phosphorylation of DDIAS on Ser676 and Thr868 by PLK1, to allow a stable and high-affinity DDIAS-TOPBP1 interaction to be established at the onset of mitosis (**Figure 4F**).

### The OB fold of DDIAS binds to single-stranded DNA (ssDNA)

Having revealed the mechanistic basis for DDIAS localization to DNA lesions, we next wished to establish its molecular function upon recruitment. While its N-terminal OB fold is not essential for DDIAS foci formation (**Figure 3B**), we considered whether it might nonetheless play a role in mediating DDIAS functions. Interestingly, the DDIAS OB fold shares homology with the C-terminal OB4 domain of RPA1, including the conserved cysteine residues that comprise a zinc-binding motif that mediates ssDNA-binding.^42^

To test whether the DDIAS OB fold binds DNA, we purified a recombinant N-terminal DDIAS fragment containing the OB fold from *E. coli* for use in electrophoretic mobility shift assays (EMSAs). Results from these experiments showed that the DDIAS OB fold could readily bind ssDNA, but showed no detectable interaction with double-stranded DNA (dsDNA; **Figure 5A**). Substituting putative zinc-binding cysteine residues with alanine (C25A/C28A) abrogated ssDNA binding, confirming that the DDIAS OB fold can engage ssDNA in a similar manner to RPA1.

**Figure 5.**
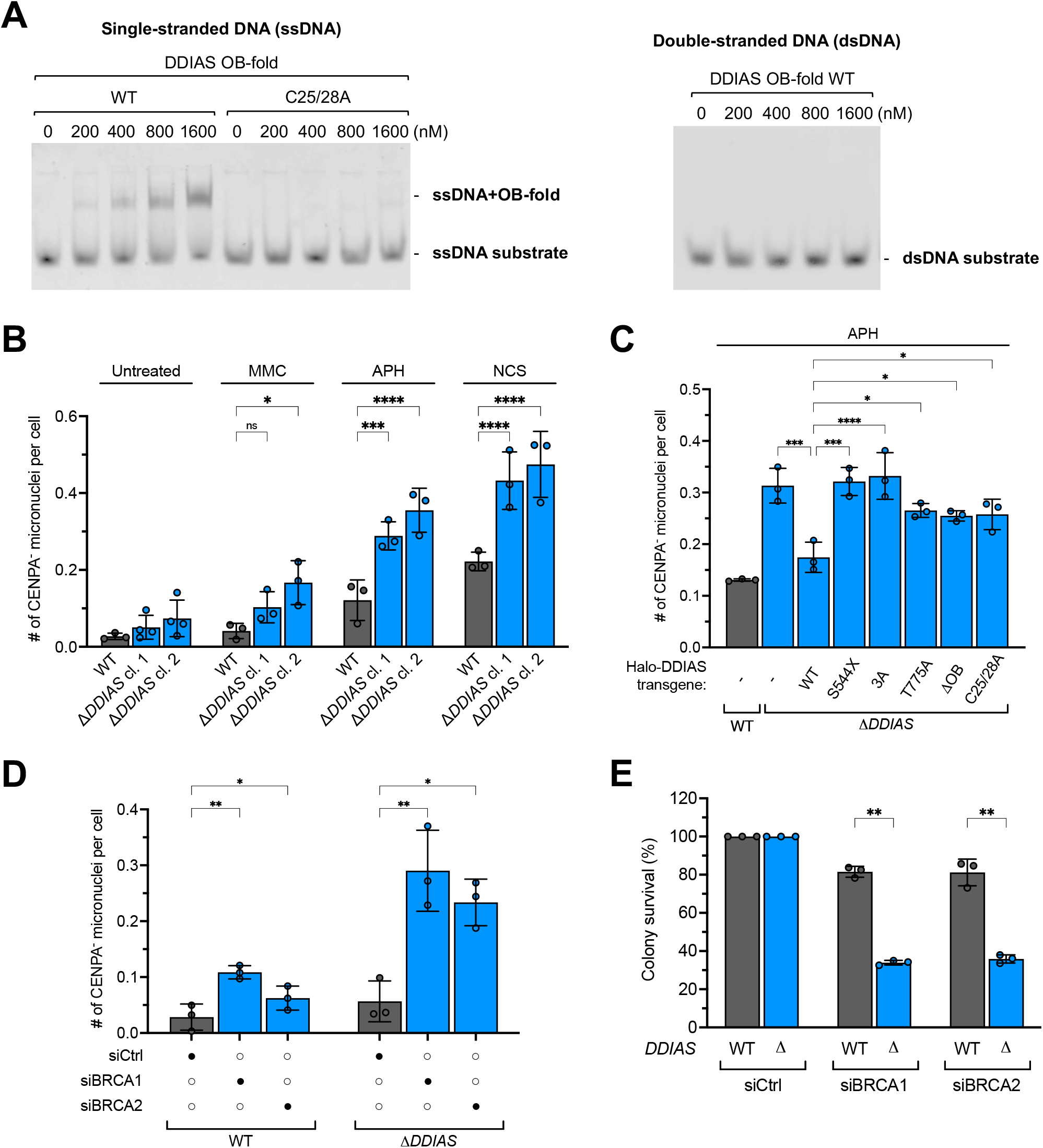
DDIAS binds ssDNA, suppresses acentric micronucleation, and promotes survival of BRCA-deficient cells. **(A)** EMSA with recombinant WT and C25A/C28A DDIAS OB fold proteins incubated at indicated concentrations with ssDNA and dsDNA substates. **(B)** Quantification of acentric (CENPA-negative) micronuclei in WT DLD-1 and two independent Δ*DDIAS* clones. Cells were mock-treated or treated with the indicated drugs for 48 hr. Error bars denote SD from n = 3 experiments. One-way ANOVA followed by Tukey’s post hoc test was used to calculate significance. **(C)** Quantification of acentric micronuclei in WT and Δ*DDIAS* DLD-1 cells expressing the indicated Halo or Halo-DDIAS proteins and treated with 400 nM APH for 48 hr. Error bars denote SD from n = 3 experiments. One-way ANOVA followed by Tukey’s post hoc test was used to calculate significance. **(D)** Quantification of acentric micronuclei in WT and Δ*DDIAS* DLD-1 cells treated with the indicated siRNAs. Error bars denote SD from n = 3 experiments. Two-way ANOVA followed by Tukey’s post hoc test was used to calculate significance. **(E)** Quantification of colony survival assays with WT and Δ*DDIAS* DLD-1 cells treated with the indicated siRNAs. Error bars denote SD from n = 3 experiments. Two-way ANOVA followed by Tukey’s post hoc test was used to calculate significance. See also **Figure S6**.

### DDIAS suppresses acentric micronucleation and its loss is synergistic with BRCA deficiency

Cells deficient in the TOPBP1-CIP2A complex display a high level of acentric micronucleation.^9,10,12^ We observed a similar phenotype in our DDIAS-deficient patient’s cells (**Figure 1E**), and in two independent Δ*DDIAS* DLD-1 clones exposed to a panel of genotoxic stress-inducing agents (**Figure 5B**). Re-expression of WT DDIAS complemented Δ*DDIAS* cells, but DDIAS mutants deficient in TOPBP1 (3A) or PLK1 binding (T775A) were unable to do so (**Figures 5C and S6A**). In addition, DDIAS mutants deficient in ssDNA binding (C25A/C28A; **Figure 5A**) or lacking the OB fold altogether, were similarly unable to suppress excess micronuclei formation, as was the truncated DDIAS protein potentially produced as a result of the patient mutation S544X (**Figures 5C, S1E and S6A**). We conclude that stable recruitment to DNA damage sites via TOPBP1 interaction is essential for DDIAS to maintain chromosome stability, as is its ability to bind ssDNA.

The TOPBP1-CIP2A complex is essential for survival of BRCA-deficient cells, presumably because the rampant chromosome breakage and micronucleation that occurs in absence of BRCA1 or BRCA2 and a functional TOPBP1-CIP2A pathway, is lethal.^10,11^ We therefore wished to assess whether DDIAS is similarly required for genome maintenance in BRCA-deficient cells. To do this, we depleted BRCA1 or BRCA2 in Δ*DDIAS* DLD-1 clones using siRNAs (**Figure S6B**), and assessed the level of acentric micronucleation in these cells. Strikingly, we observed a very large induction of micronuclei in cells lacking DDIAS and either BRCA1 or BRCA2 (**Figure 5D**). Consistent with this, Δ*DDIAS* cells were also significantly compromised in their ability to survive when depleted of BRCA1 or BRCA2 (**Figure 5E**); similar results were observed in reciprocal experiments where we depleted DDIAS using two different siRNAs in Δ*BRCA2* DLD-1 cells (**Figures S6C and S6D**). Thus, we conclude that the cytogenetic burden imposed by homologous recombination deficiency becomes lethal when DDIAS is lost, in line with its role as a downstream factor in the TOPBP1-CIP2A pathway.

## DISCUSSION

In this study, we reveal DDIAS as a novel member of the TOPBP1-CIP2A complex in a mitotic DNA damage response that safeguards chromosome integrity, particularly in BRCA-deficient cells. Our data support a model in which DDIAS acts downstream of TOPBP1 and CIP2A recruitment, to endow the complex with ssDNA-binding activity via its N-terminal OB fold in a manner that is tightly controlled by CDK1 and PLK1 kinases to ensure its role is specific to mitosis (**Figure 6**).

**Figure 6.**
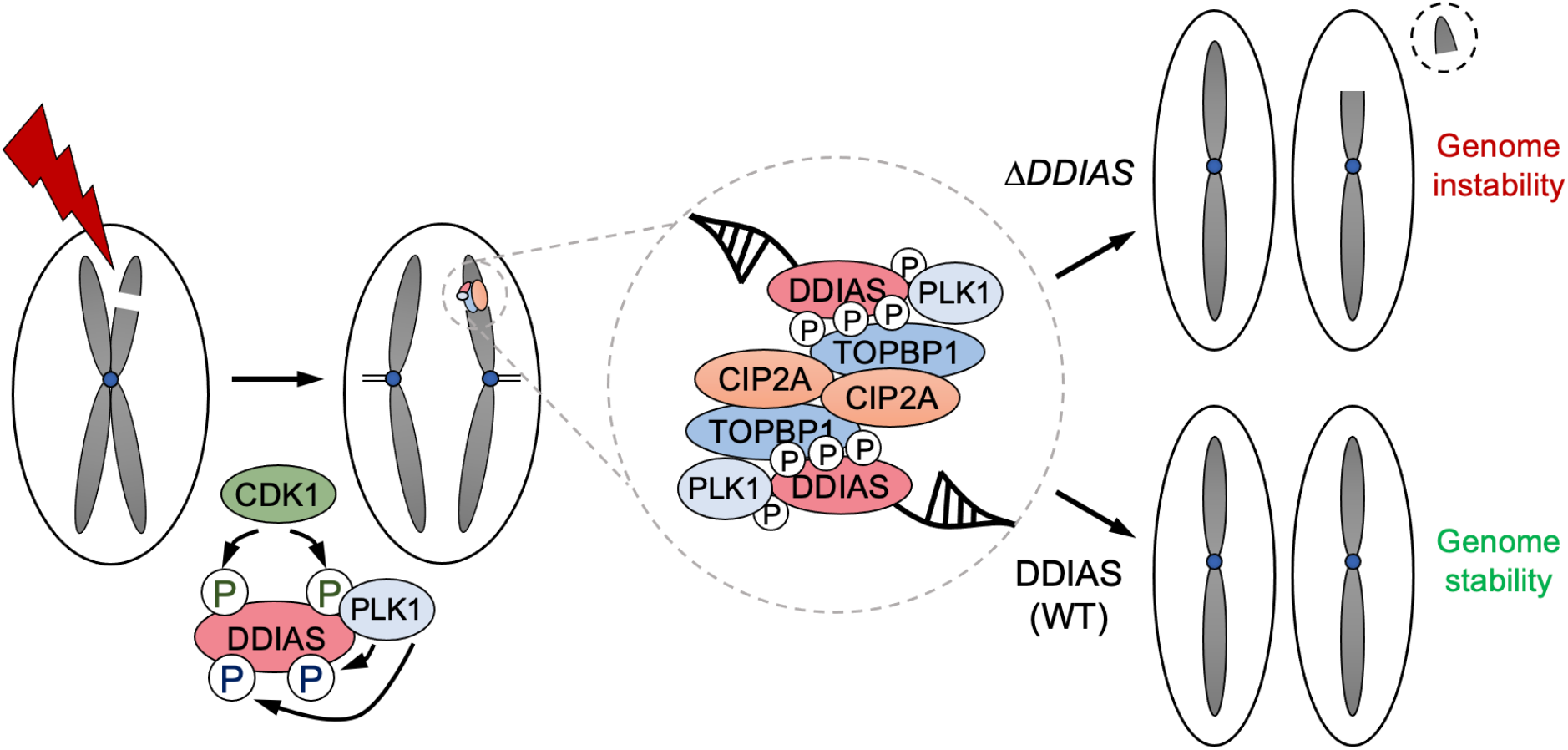
Model for the role of DDIAS in the TOPBP1-CIP2A pathway of genome stability maintenance in mitosis.

Currently, it is not clear how TOPBP1-CIP2A-DDIAS complexes function to maintain chromosomal stability during mitosis. We could find no evidence for a role of DDIAS in mitotic DNA synthesis (MiDAS), MMEJ or other DSB repair pathway during mitosis (**Figures S6E and S6F**), despite previous reports of TOPBP1 and/or CIP2A in these processes.^20,25,28,30^ It is possible that DDIAS acts either to protect chromosomes containing ssDNA gaps from breakage by spindle forces or aberrant enzymatic processing, or to bridge under-replicated regions during mitosis after nucleolytic cleavage or DNA unwinding has already occurred. This would allow the physical tethering of acentric fragments to their centromere-containing chromosome counterparts during cell division until the following G1 phase in daughter cells, when canonical DNA repair pathways are reactivated.^4^ Both models are supported by the observation that mutation of the DDIAS OB fold or its TOPBP1-binding sites have similar consequences to loss of DDIAS altogether, suggesting a crucial role for ssDNA-binding by DDIAS as part of the TOPBP1-CIP2A complex. Either model could also plausibly explain the synthetic lethality between BRCA loss and deficiency in the TOPBP1-CIP2A-DDIAS complex, given that BRCA-deficient cells are known to accumulate under-replicated regions, ssDNA gaps and DSBs during replication.^43,44^

In conclusion, our identification of DDIAS as a ssDNA-binding component of the TOPBP1-CIP2A complex provides insight into its role in mitosis and reveals a tractable pathway for therapeutic intervention in BRCA-deficient and potentially other cancers in combination with PARP, POLQ or ATR inhibitors. The role of the TOPBP1-CIP2A-DDIAS pathway in human neurodevelopment also warrants further investigation; future studies with a larger cohort of patients will be required to establish this. Notably, neural progenitors undergo extraordinarily rapid divisions during corticogenesis, with cell cycles as short as 8 hours in initial stages of differentiation;^45^ they may thus be more likely to enter mitosis with under-replicated regions and other replication-derived DNA lesions than most other cell types, and thus be highly reliant on TOPBP1-CIP2A-DDIAS to maintain chromosome stability.

## ACKNOWLEDGMENTS

We are particularly grateful to the patient and their family for their participation in this study. We thank Thanos Halazonetis, Philip Hublitz, Luke Lavis, Tobias Meyer, Manuel Stucki, Madalena Tarsounas, Didier Trono and Feng Zhang for providing reagents. We thank Ian Hickson and Jiri Lukas for helpful discussions, and Jonathan Levy and Daniel Durocher for sharing and discussing results prior to publication. Imaging was carried out with help from the University of Copenhagen’s Core Facility for Integrated Microscopy (CFIM). Mass spectrometry was carried out by the University of St Andrews Biomedical Sciences Research Complex Mass Spectrometry & Proteomics Core Facility, or the University of Copenhagen’s Protein Research Infrastructure, supported by the Novo Nordisk Foundation (NNF19SA0059305). We are grateful to Jan Bisenberger for providing technical support. The Blackford laboratory was supported by a Cancer Research UK (CRUK) Senior Cancer Research Fellowship to A.N.B. (RCCSCF-May23/100001), a Novo Nordisk Foundation Hallas-Møller Ascending Investigator award to A.N.B. (NNF23OC0082165), a Japan Society for the Promotion of Science Overseas Research Fellowship to K.T. (202260198), a Uehara Memorial Foundation Postdoctoral Fellowship to K.T. (202331053), a Lundbeck Foundation Fellowship to K.T. (R483-2024-1673), and a Novo Nordisk Foundation BRIDGE Fellowship to L.L. (NNF23SA0087869). The Ochs laboratory was supported by a Novo Nordisk Foundation grant (NNF23OC0084734) and an Independent Research Fund Denmark Inge Lehmann grant (10.46540/4305-00006B). The Stewart laboratory was supported by a CRUK Programme Grant (C17183/A23303), a BBSRC project grant (BB/Y010078/1), and a Birmingham Health Partners (BHP) Starter Fellowship to C.S.S. The Miller laboratory was supported by the Danish National Research Foundation (DNRF115), the Carlsberg Foundation (CF21-0571), and a Novo Nordisk Foundation Hallas-Møller Emerging Investigator award to T.C.R.M. (NNF22OC0073571). A.F. received funding from the International Centre for Genetic Engineering and Biotechnology (ICGEB; CRP/PAK21-05_EC).

## AUTHOR CONTRIBUTIONS

Conceptualization: A.N.B. and F.O. Investigation: K.T., L.L., A.T., I.I.G., C.S.S., M.K., A.W.O., J.B., G.S.S., F.O. and A.N.B. Resources: S.E.J., S.S.J., A.A., M.R., M.T., Z.I. and A.F. Supervision: A.N.B., F.O., G.S.S. and T.C.R.M. Writing – original draft: A.N.B. and K.T. Writing – review & editing: K.T., L.L., I.I.G., A.W.O., S.E.J., J.B., Z.I., T.C.R.M., G.S.S., F.O. and A.N.B.

## STAR METHODS

### Research subject inclusion and ethics

Informed consent was obtained from the participating family to take clinical samples, perform DNA and data analyses, and to publish clinical information and photographs in accordance with local approval regulations and in compliance with Declaration of Helsinki principles. This study was approved by the West Midlands, Coventry and Warwickshire Research Ethics Committee (REC: 20/WM/0098) and the Ethics Review Committee of the Aga Khan University Hospital (ERC: 2021-6514-19346).

### Clinical synopsis and molecular genetics

DDIAS-P1 is a five-year-old male child of Pakistani origin from a consanguineous marriage. The proband was born at full term but started having frequent generalized clonic seizures one week after birth, that could be controlled with medication. He presented with a thin wasted appearance, global developmental delay, microcephaly, short stature, intellectual disability, limited speech, generalized spasticity, brisk reflexes in all four limbs, horizontal nystagmus, high arched palate, bilateral clinodactyly and skin hypopigmentation. The proband was unable to sit, stand or hold his neck up unaided. His vision and hearing were unaffected. Recurrent chest infections were also reported. Nuclear magnetic resonance imaging (MRI) of the affected child’s brain revealed periventricular encephalomalacia, abnormal signals in the pulvinar nuclei, ventriculomegaly, thinning of the corpus callosum and mild diffuse cerebral atrophic changes including suspected perinatal bleeding. Whole exome sequencing (WES) was carried out to identify the underlying cause of the affected child’s neurological abnormalities. This analysis identified a likely pathogenic homozygous nonsense variant in *DDIAS* (NM_145018.4:c.1631C>A; p.Ser544Ter), with a CADD (v1.6) score of 33, which was identified in gnomAD (v4.1.0) at an allele frequency of 1.24 x 10^-5^ with no homozygotes present. The variant was present in a heterozygous state in the proband’s parents and his unaffected sister. No likely causative variants in other genes were evident in the WES dataset.

### Cell lines and culture conditions

All cells were grown in humidified incubators supplied with 5% CO_2_ and maintained at 37 °C, with regular testing for mycoplasma contamination using a MycoAlert Mycoplasma Detection Kit (Lonza). RPE-1 FRT/TR Δ*TP53* (referred to as RPE-1 throughout this study) cells have been described previously.^21^ RPE-1 Δ*CIP2A* and their parental cells were gifts from Manuel Stucki.^12^ U2OS cells were obtained from the American Type Culture Collection (ATCC). DLD-1 cells and their Δ*BRCA2* derivatives were gifts from Madalena Tarsounas.^46^ RPE-1, U2OS and DLD-1 cells were cultured in Dulbecco’s modified Eagle’s medium (DMEM) containing GlutaMAX (Thermo Fisher Scientific), supplemented with 10% fetal calf serum (FCS; Thermo Fisher Scientific) and 100 U/mL penicillin/100 µg/mL streptomycin (Thermo Fisher Scientific). 293FT cells were obtained from Thermo Fisher Scientific and cultured in DMEM containing GlutaMAX, supplemented with 10% FCS, 1% MEM non-essential amino acids (Thermo Fisher Scientific) and 500 μg/mL Geneticin (G418; Thermo Fisher Scientific). Unless otherwise indicated, doxycycline was added to a final concentration of 100 ng/mL. Patient-derived primary dermal fibroblasts were grown from skin-punch biopsies and maintained in DMEM supplemented with 20% FCS, L-glutamine and penicillin-streptomycin. Patient-derived lymphoblastoid cell lines were generated from peripheral blood samples, transformed with Epstein-Barr virus, and were maintained in Roswell Park Memorial Institute (RPMI) 1640 medium (Thermo Fisher Scientific), supplemented with 10% FCS, glutamine and penicillin-streptomycin.

### Micronuclei quantification

Where indicated, cells were transfected with siRNA 24 hr prior to plating on glass coverslips, and then mock-treated or treated with APH, MMC or NCS for 48 hr. Cells were fixed in 4% paraformaldehyde in phosphate-buffered saline (PBS) for 10 min at room temperature, washed twice with PBS, permeabilized and blocked in blocking buffer (3% BSA, 0.5% Triton X-100 in PBS) for 30 min at room temperature. Coverslips were washed in PBS, then incubated with anti-CENPA primary antibody (ADI-KAM-CC006-E, Enzo Life Sciences, 1/1000) in blocking buffer for 1 hr at room temperature. Cells were washed three times in PBS-T (0.2% Tween 20 in PBS), before incubation with goat anti-mouse IgG Alexa Fluor Plus 488 secondary antibody (A32723, Thermo Fisher Scientific, 1/1000) in blocking buffer containing 0.5 μg/mL DAPI in the dark for 1 hr at room temperature. Cells were washed three times in PBS-T and once in water, before mounting in 5 μL of a 10% Mowiol 4-88 (Merck), 25% glycerol, 100 mM Tris-HCl, pH 8.5 solution. Images were acquired using an Olympus BX63 microscope with a UPlanFL N 40x/0.75 objective. Micronuclei were scored manually using Fiji.^47^

### Analyses of chromosomal breakage

Patient-derived cells were blocked in mitosis by addition of 200 ng/mL KaryoMAX Colcemid Solution in PBS (Thermo Fisher Scientific) for 3 hr. After harvesting by trypsinization, cells were resuspended in hypotonic buffer (10 mM KCl, 15% FCS) and incubated at 37 °C for 30 min, followed by fixation by addition of 3:1 ethanol/acetic acid solution. Cells were dropped onto acetic-acid-humidified slides, stained for 15 min in 5% modified Giemsa stain (Merck) in water, and washed for 5 min in water.

### Plasmids and cloning

All plasmids were prepared using NucleoSpin Plasmid Mini (Macherey-Nagel) or GenElute HP Endotoxin-Free Plasmid Maxiprep (Merck) kits according to the manufacturer’s instructions, and verified by Sanger or Oxford Nanopore Technologies sequencing (Eurofins Genomics). Point mutations were introduced by site-directed mutagenesis using QuikChange (Agilent Technologies). HaloTag-fused human DDIAS cDNA was synthesized and cloned into pTwist-CMV (Twist Biosciences). For lentiviral transductions, Halo-DDIAS cDNAs were amplified by PCR from pTwist-CMV and inserted into a modified TLCV2 vector^21^ to replace the Cas9 cassette using In-Fusion cloning (Takara Bio) according to the manufacturer’s instructions. The pIRESneo2 plasmid encoding GFP-TOPBP1 was a gift from Thanos Halazonetis.^48^ GFP-TOPBP1 mutant derivatives were described previously,^9,23,24^ except for the BRCT5+7 double mutant containing K704A and K1317A mutations, which was generated during the course of this study. For lentiviral transductions, TOPBP1 cDNA from the pIRESneo2 plasmid encoding GFP-TOPBP1 was inserted into a pLenti PGK Neo DEST vector expressing mClover3-BLM and FLAG-RMI1^49^ using In-Fusion cloning to replace the BLM and FLAG-RMI1 sequences, producing a lentiviral vector expressing mClover3-TOPBP1.

### Lentiviral transductions

To assemble lentiviruses for exogenous protein expression, 293FT cells were transfected with vectors encoding mClover3-TOPBP1 or Halo-DDIAS proteins, along with pHDM-tat1b, pHDM-G, pRC/CMV-rev1b and pHDM-Hgpm2 lentiviral assembly plasmids.^50^ Viral supernatants were harvested at 24 and 48 hr, pooled, passed through a 0.45-μm filter, and stored at -80 °C. For lentiviral infections, RPE-1, DLD-1, U2OS or patient-derived cells were incubated in viral media containing 0.8 μg/mL polybrene for 24 hr, washed in medium twice and incubated with fresh medium for 24 hr, before selection in fresh medium containing 500 μg/mL G418 or 1 μg/mL puromycin (Thermo Fisher Scientific) for up to 7 days. G418-resistant U2OS cells were sorted for mClover3-TOPBP1 expression. All stable cell lines were verified by western blotting.

Patient-derived fibroblasts were immortalized with a lentivirus expressing human telomerase reverse transcriptase (hTERT) that was generated by transfecting 293FT cells with the following plasmids: pLV-hTERT-IRES-hygro (a gift from Tobias Meyer; Addgene plasmid # 85140), psPAX2 (a gift from Didier Trono; Addgene plasmid # 12260) and pMD2.G (a gift from Didier Trono; Addgene plasmid # 12259). Selection was performed using 70 μg/mL hygromycin B (Thermo Fisher Scientific).

### SDS-PAGE and western blotting

SDS-PAGE and western blotting were performed using 7% Tris-Bicine gels with SE400 and TE42 systems from Hoefer, or Bolt 4%–12% Bis-Tris Plus gels from Thermo Fisher Scientific. The following antibodies were used at the indicated dilutions: BLM (A300-110A, Bethyl Laboratories, 1/2000), BRCA1 (sc-6954, Santa Cruz Biotechnology, 1/200), BRCA2 (OP95, Merck, 1/2000), CIP2A (14805, Cell Signaling Technology, 1/1000), FANCJ (4578, Cell Signaling Technology, 1/500), GFP (11814460001, Roche, 1/5000), H2AX (NB100-383, Novus Biologicals, 1/5000), HaloTag (G9211, Promega, 1/1000), PLK1 (05-844, Merck, 1/4000), RAD9 (sc-8324, Santa Cruz Biotechnology, 1/1000), RPA2 (ab10359, Abcam, 1/10,000), TOPBP1 (A300-111A, Bethyl Laboratories, 1/1000).

### Generation of *DDIAS* knockout cells

To knock out *DDIAS* in RPE-1 and DLD-1 cells, small guide RNAs (sgRNAs) were designed targeting the following sequences: sgDDIAS-A1, 5’-TAGTTACCTGCCTAACCATC-3’; sgDDIAS-A2, 5’-TAGGAGTTCTGTACATGTGA-3’; sgDDIAS-B1, 5’-CAGTAAAGCCTGCAATACCT-3’; sgDDIAS-B2, 5’-AGAACTTGGCTTACAAGCTA-3’. sgDDIAS-A1 and sgDDIAS-B1 were cloned into pSpCas9(BB)-2A-GFP (PX458; a gift from Feng Zhang;^51^ Addgene plasmid # 48138). sgDDIAS-A2 and sgDDIAS-B2 were cloned into a modified PX458 in which GFP was replaced with mRuby2 (a gift from Philip Hublitz^52^). Plasmids were transfected into cells using Lipofectamine 3000 (Thermo Fisher Scientific), according to the manufacturer’s instructions. Cells were left to recover for 3 days before sorting GFP- and mRuby2-positive cells. Colonies were grown from single cells and screened by PCR.

### Immunofluorescence and confocal microscopy

Cells were grown on 12-mm-wide, 0.13-0.16-mm-thick glass coverslips (VWR; cleaned in 96% ethanol, dried, and autoclaved), and irradiated when appropriate using a Faxitron CP-160 X-ray generator. Where indicated, the POLQ inhibitor ART558^53^ as added at 10 µM concentration. To quantify DDIAS or CIP2A foci in mitosis, cells were treated with 400 nM APH for 8 hr followed by addition of 100 ng/mL nocodazole for 16 hr to enrich for mitotic cells, unless otherwise indicated. Cells were fixed in 4% formaldehyde and 0.1% Triton X-100 in PBS for 15 min, washed twice with PBS, permeabilized in 0.5% Triton X-100 in PBS for 5 min, and blocked in blocking buffer for 30 min, all at room temperature. Coverslips were washed in PBS-T, then incubated at room temperature for 1 hr with one or more of the following primary antibodies as indicated, diluted in antibody buffer (DMEM supplemented with 10% FCS and 0.05% sodium azide; filtered): CIP2A (Abcam, ab99518, 1/400), γH2AX (Merck, 05-636, 1/1000), H3-pS10 (mouse, Abcam, ab14955, 1/1000 or rabbit, Merck, 06-570, 1/1000), TOPBP1 (Santa Cruz Biotechnology, sc-271043, 1/200). Cells were then washed three times in PBS-T, before incubation in the dark for 1 hr at room temperature with the following secondary antibodies as appropriate, diluted in antibody buffer containing 0.5 μg/mL DAPI: goat anti-mouse IgG Alexa Fluor Plus 488 (A32723, 1/1000), goat anti-rabbit IgG Alexa Fluor Plus 488 (A32731, 1/1000), goat anti-mouse Alexa Fluor Plus 555 (A32727, 1/1000), goat anti-rabbit Alexa Fluor Plus 555 (A32732, 1/1000), goat anti-mouse Alexa Fluor Plus 647 (A32728, 1/1000), goat anti-rabbit Alexa Fluor Plus 647 (A32733, 1/1000); all from Thermo Fisher Scientific. Cells were washed three times in PBS-T and once in water, before mounting in 5 μL of a 10% Mowiol 4-88 (Merck), 25% glycerol, 100 mM Tris-HCl, pH 8.5 solution or SlowFade Diamond Antifade mountant (Thermo Fisher Scientific). For detection of Halo-DDIAS, labeling was carried out by incubating cells with the HaloTag ligand dye JFX554 (a gift from Luke Lavis) at 100 nM for 1.5 hr. Cells were washed twice in medium, and incubated with fresh medium for 15 min as an exit wash to remove unbound dye before fixation. Where indicated, EdU was labeled with Alexa Fluor 647 using a Click-iT EdU Cell Proliferation Kit for Imaging (Thermo Fisher Scientific) according to the manufacturer’s instructions prior to antibody incubations. Confocal images were acquired using a ZEISS LSM 780 inverted microscope equipped with a 63x/1.4 NA Plan-Apochromat objective. DAPI was detected using a 405-nm diode laser, Alexa 488, GFP and mClover3 were detected with the 488-nm line of an argon laser, Alexa 555 and JFX554 were detected with the 543-nm line of a HeNe laser and Alexa 647 was detected with the 633-nm line of a HeNe laser. Z stacks of entire cells were acquired using optimized step sizes and processed by ZEN Black 2012 software. Display of images was adjusted for intensity for optimal display of structures of interest.

### Quantitative Image-Based Cytometry (QIBC)

QIBC experiments were performed on an Olympus scanR inverted microscope system equipped with wide-field optics, a 20x/0.8 NA dry objective, fast excitation and emission filter wheel devices for DAPI, FITC, Cy3, and Cy5 wavelengths, an MT20 illumination system, and a digital monochrome Hamamatsu ORCA-Flash 4.0 digital camera, yielding a spatial resolution of 320 nm per pixel at 20x and binning 1. Acquisition settings for each channel were optimized for optimal signal over noise ratios and non-saturated conditions. Identical acquisition settings were applied across all samples within one experiment and between repeats. Images were processed and analyzed using scanR acquisition and image analysis software (Olympus, version 3.5.0). Firstly, mitotic cells were segmented based on H3-pS10 staining using an intensity threshold mask. This mask was then applied to quantify pixel intensities across other fluorescence channels for each identified mitotic cell. Foci were then segmented using intensity-based thresholding and size-exclusion. Following segmentation, a range of parameters was extracted for both whole mitotic cells and foci. These included direct measurements (e.g. mean and total intensities, area, circularity, foci count, and foci intensities) as well as derived metrics (e.g. sum of foci intensity per mitotic cell). In addition, the segmented objects were gated for circularity and area to exclude under- and over-segmented objects. The resulting data were exported in tabular format and further analyzed using Spotfire (TIBCO Software, version 14.0.1). For DDIAS overexpression experiments, cells were gated by DDIAS total intensity per mitotic cell and only the lowest 20% expressing cells were used for further analysis, to prevent segmentation artefacts caused by protein overexpression. Within each experiment, comparisons were made between conditions using similar cell counts. For visualization, low y-axis jittering was applied (random displacement of objects along the y-axis) to make overlapping markers visible.

### Peptide pulldowns

Peptides were synthesized by Genosphere Technologies and biotinylated on the N terminus with the following sequences: SGSG-EGSYDASADLFDD (DDIAS-S676), SGSG-EGSYDA-[pSer]-ADLFDD (DDIAS-pS676), SGSG-SQDFVPCSQSTPIS (DDIAS-T775), SGSG-SQDFVPCSQS-[pThr]-PIS (DDIAS-pT775), SGSG-ETDSDEWVPPTTQK (DDIAS-T868), SGSG-ETDSDEWVPP-[pThr]-TQK (DDIAS-pT868), SGSG-ETRSAWSPELFS (DDIAS-S993), SGSG-ETRSAW-[pSer]-PELFS (DDIAS-pS993). Peptides were bound to streptavidin-coupled Dynabeads M-280 (Thermo Fisher Scientific). Lysates were prepared by washing 293FT cells in PBS and lysing them in nuclease buffer (100 mM NaCl, 0.2% Igepal CA-630, 1 mM MgCl_2_, 10% glycerol, 5 mM NaF, 50 mM Tris-HCl, pH 7.5) supplemented with cOmplete EDTA-free protease inhibitor cocktail (Roche) and 25 U/mL SuperNuclease (Sino Biological). After nuclease digestion, NaCl concentration was adjusted to 200 mM and 2 mM EDTA was added, and lysates were cleared by centrifugation. Dynabead-conjugated peptides were incubated with clarified extracts with end-to-end mixing at 4 °C for 2 hr. Beads were then washed five times with wash buffer (nuclease buffer supplemented with 2 mM EDTA and NaCl concentration adjusted to 200 mM). For western blotting, proteins were eluted in 2X SDS sample buffer (4% SDS, 20% glycerol, 50 mM TCEP, 0.002% bromophenol blue, 125 mM Tris-HCl, pH 6.8). For mass spectrometry, beads were washed in Tris-buffered saline (TBS) and stored at -20 °C.

### Mass spectrometry

Proteins for mass spectrometry were subjected to on-bead digestion with trypsin. Peptides were subjected to LC-MS/MS using an UltiMate 3000 rapid separation LC (RSLC; Thermo Fisher Scientific) coupled to an Orbitrap Fusion Lumos mass spectrometer with a field-asymmetric ion mobility spectrometry (FAIMS) interface (Thermo Fisher Scientific). Peptides were injected onto a PepMap 100 C18 (5 μm, 100 μm x 2 cm) reverse-phase trap for pre-concentration and desalted with 0.05% trifluoroacetic acid in water, at 5 μL/min for 10 min. The peptide trap was then switched into line with the analytical column (EASY-Spray PepMap RSLC C18 2 μm, 15 cm x 75 μm internal diameter). Peptides were eluted from the column using a linear solvent gradient (mobile phase A: 0.1% formic acid in water, mobile phase B: 80% acetonitrile, 0.1% formic acid, 20% water) using the following gradient: linear 2–35% of buffer B over 44 min, sharp increase to 95% buffer B within 0.5 min, isocratic 95% of buffer B for 4.5 min, sharp decrease to 2% buffer B within 1 min and isocratic 2% buffer B for 5 min. The FAIMS interface was set to -45 V and - 65 V at standard resolution. The mass spectrometer was operated in data-dependent acquisition positive ion mode with a cycle time of 1.5 s. The Orbitrap was selected as the MS1 detector at a resolution of 60,000 with a scan range of from m/z 375 to 1500. Peptides with charge states 2 to 5 were selected for fragmentation in the ion trap using higher-energy collisional dissociation (HCD) as collision energy. Raw files were processed with MaxQuant version 2.6.7.0 with integrated Andromeda search engine against the human UniProt database (UP000005640_9606). Carbamidomethylation of cysteine was set as a fixed modification, while oxidation of methionine and protein N-acetylation were considered variable modifications. Trypsin was selected as an enzyme with a maximum of two missed cleavages. Search results were filtered with a false discovery rate of 0.01. Label-free quantification for unique + razor peptides and the match between run option were activated in the MaxQuant software. Further analysis was performed in Perseus or Proteomelit software and included data filtering, imputation, normalization and statistics (t-tests with FDR correction).

### Fluorescence polarization

cDNAs encoding residues 548-741 (BRCT4+5) or 1263-1495 (BRCT7+8) of human TOPBP1 were codon-optimised for *E. coli* expression, synthesized (Thermo Fisher Scientific), and cloned into the modified pET-17b vector pAWO-His-SUMO-3C^54^ using Gibson assembly. Proteins were expressed in *E. coli* BL21(DE3) by induction with 0.2 mM ITPG at OD_600_ of 1.5 at 20 °C for 16 hr. After induction, cells were harvested by centrifugation and pellets were frozen at -20 °C. Pellets were resuspended in buffer A (250 mM NaCl, 10 mM imidazole, 0.5 mM TCEP, 50 mM HEPES-NaOH, pH 7.5), supplemented with protease inhibitors (Roche), and lysed by sonication. After centrifugation, supernatants were applied to a gravity column containing 1 mL of TALON affinity capture resin (Takara Bio). After successive washes with buffer B (250 mM NaCl, 10 mM imidazole, 0.5 mM TCEP, 20 mM HEPES-NaOH, pH 7.5), protein was eluted by application of buffer C (250 mM NaCl, 300 mM imidazole, 0.5 mM TCEP, 20 mM HEPES-NaOH, pH 7.5). Selected fractions were pooled after analysis by SDS-PAGE, and then applied to a HiLoad 16/600 Superdex 75 size exclusion chromatography column (Cytiva) pre-equilibrated in Buffer D (200 mM NaCl, 0.5 mM TCEP, 20 mM HEPES-NaOH, pH 7.5). Analysis by SDS-PAGE identified fractions containing purified protein, which were then pooled, concentrated, flash-frozen and stored at -80°C until required. Peptides were synthesized by Genosphere Technologies and labelled with 5-carboxy-fluoroscein on the N terminus with the following sequences: GSG-EGSYDA-[pSer]-ADLFDD (DDIAS-pS676), GSG-ETDSDEWVPP-[pThr]-TQK (DDIAS-pT868), GSG-ETRSAW-[pSer]-PELFS (DDIAS-pS993). Fluorescence polarization experiments were performed in 200 mM NaCl, 0.5 mM TCEP, 0.05% Igepal CA-630, 20 mM HEPES-NaOH, pH 7.5, and measured using a CLARIOstar multi-mode microplate reader (BMG Labtech). Data were analyzed using GraphPad Prism (version 10.4.1), by non-linear fitting with a one-site specific binding model. Data represent the mean of three experiments, and error bars represent one standard deviation.

### Immunoprecipitations

Plasmids were transfected into 293FT cells using Lipofectamine 2000 (Thermo Fisher Scientific) according to the manufacturer’s instructions. Unless otherwise stated, 100 ng/mL nocodazole was added for 16 hr to enrich for mitotic cells. For preparation of lysates for immunoprecipitations, cells were washed in PBS, and lysed in nuclease buffer supplemented with cOmplete EDTA-free protease inhibitor cocktail and 25 U/mL SuperNuclease. In experiments for phospho-proteomics, buffer was additionally supplemented with PhosStop phosphatase inhibitor cocktail (Roche). After nuclease digestion, NaCl and EDTA concentrations were adjusted to 200 mM and 2 mM, respectively, and lysates were cleared by centrifugation. Lysates were then incubated with GFP-Trap or Halo-Trap magnetic agarose beads (Proteintech) for 2 hr with end-to-end mixing at 4 °C. Immunoglobulin-antigen complexes were washed five times in wash buffer, before elution in 2X SDS sample buffer for SDS-PAGE.

### Electrophoretic mobility shift assay (EMSA)

cDNAs encoding the DDIAS OB fold (residues 1-168) were amplified by PCR from pTwist-CMV expressing Halo-DDIAS (WT and C25A/C28A), and inserted into pET-28a(+) for bacterial expression. Proteins were expressed in *E. coli* BL21(DE3) by induction with 0.5 mM ITPG at OD_600_ of 0.6 at 37 °C for 4 hr. After induction, cells were harvested by centrifugation at 4 °C. Cell pellets were resuspended in lysis buffer (500 mM NaCl, 10% glycerol, 0.5 mM TCEP, 0.5 mg/mL lysozyme, 50 mM Tris-HCl, pH 8.0) and disrupted by sonication. After centrifugation, pellets were resuspended in denaturing buffer (6 M guanidine hydrochloride, 150 mM NaCl, 0.5 mM TCEP, 50 mM Tris-HCl, pH 8.0) and incubated overnight at 4 °C. Samples were clarified by centrifugation before being loaded onto a 1 mL Ni-NTA resin. After washing with denaturing buffer, proteins were eluted in denaturing buffer supplemented with 250 mM imidazole. For refolding, eluted proteins were transferred to a dialysis membrane (MWCO ∼10 kDa) and dialyzed against denaturing buffer with decreasing concentrations of guanidine hydrochloride every 2 hr, before a final dialysis step overnight in 150 mM NaCl, 0.5 mM TCEP, 50 mM Tris-HCl, pH 8.0. After dialysis, samples were centrifuged and analyzed by SDS-PAGE. For preparation of DNA substrates, chemically synthesized oligonucleotides (5’-Cy5-GATGAGATTGAGGCTGGCCTGCAGGCATGCAAGCACCTGG-3’ and 5’-CCAGGTGCTTGCATGCCTGCAGGCCAGCCTCAATCTCATC-3’) were resuspended in TE (1 mM EDTA, 10 mM Tris-HCl, pH 7.5). dsDNA was prepared by annealing oligonucleotides in a 1:1 ratio at 95 °C for 5 min, with slow cooling at 1 °C per minute to 20 °C. For ssDNA, only the Cy5-labelled oligonucleotide was used. Protein-DNA binding reactions for EMSA were performed in binding buffer (50 mM KCl, 0.5 mM TCEP, 0.5 mg/mL BSA, 5% glycerol, 20 mM Tris-HCl, pH 8.0) with 10 nM DNA substrate and varying amounts of protein at room temperature for 30 min in a 20 µl reaction volume. Samples were loaded onto a 6% native PAGE gel in Tris-borate-EDTA (TBE) buffer. Fluorescently labelled DNA was detected using a Typhoon FLA 9500 (Cytiva).

### RNA interference

siRNAs were transfected using Lipofectamine RNAiMAX (Thermo Fisher Scientific) according to the manufacturer’s instructions. The following sequences were used: siCtrl (targeting firefly luciferase), 5’-CGUACGCGGAAUACUUCGA-3’; siDDIAS #1, 5’-GAUACAACUCAGAAUCUAU-3’; siDDIAS #2, 5’-GAUUCAUGGAGCCUUGUUU-3’. siBRCA1 and siBRCA2 are siPOOLs of 30 siRNAs (siTOOLs Biotech), which reduce off-target effects by including very small concentrations of each individual siRNA while still achieving efficient knockdowns with 10-fold less total siRNA concentration compared to single siRNA transfections.^55^

### Colony survival assays

Where indicated, cells were transfected with siRNA 24 hr prior to plating at low densities. Cells were left for up to 14 days to allow colonies to develop. Colonies were washed in PBS, stained in 0.1% Coomassie Brilliant Blue R 250, 7% acetic acid, 50% methanol at room temperature for 30 min, and washed in water before counting.

---

**Figure S1.**
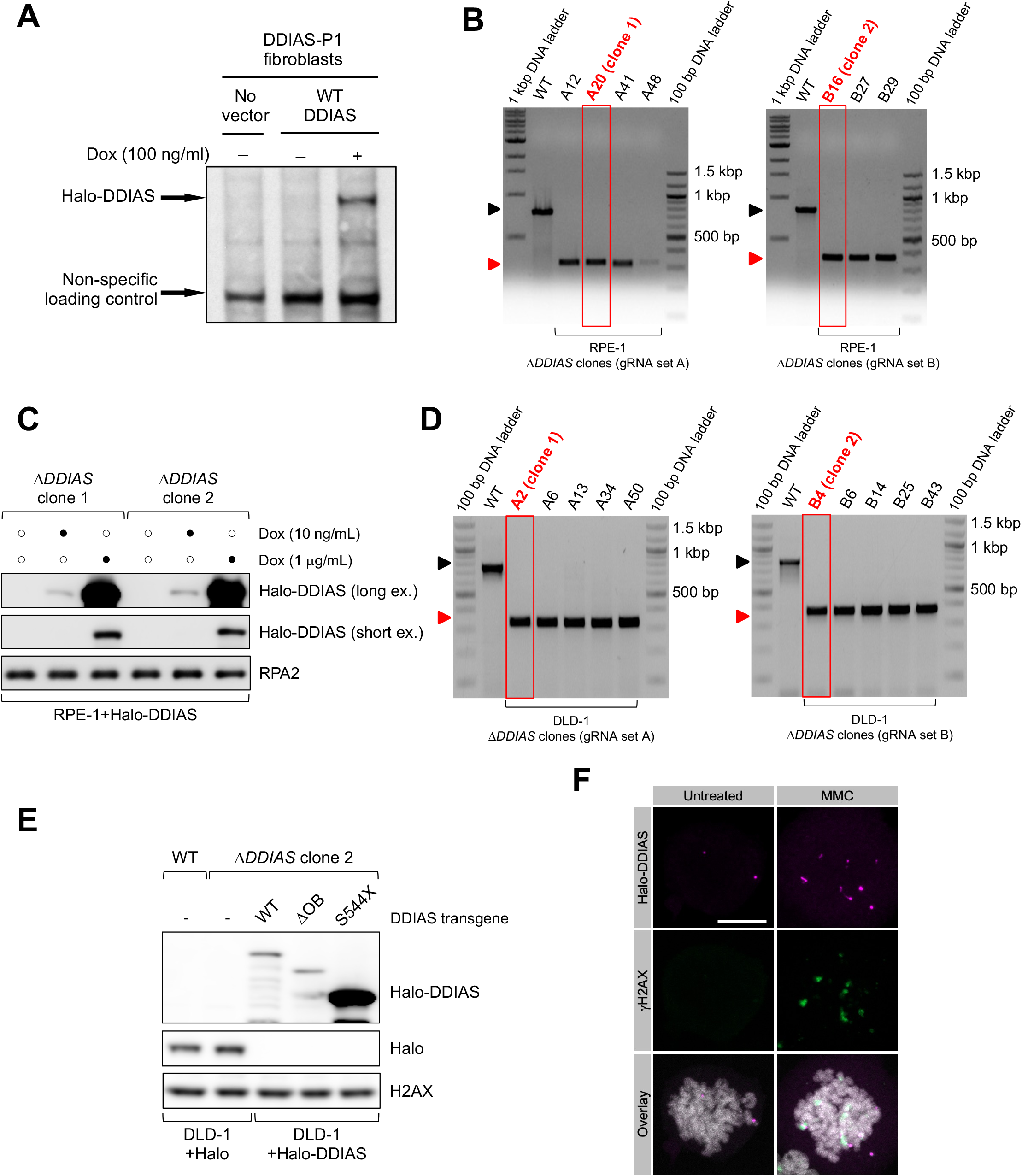
Generation of DDIAS knockout RPE-1 and DLD-1 cells and complementation with doxycycline-inducible Halo or Halo-DDIAS proteins, related to Figures 1 and 2. **(A)** Western blotting of lysates from patient-derived fibroblasts expressing Halo-tagged DDIAS in a doxycycline-inducible manner. **(B)** Agarose gel electrophoresis of PCR results showing homozygous knockout of the *DDIAS* gene in five independent clones in RPE-1 cells using two independent sets of gRNAs (gRNA set A or B). Red boxes indicate the clones selected for further analysis in this study. **(C)** Western blotting of lysates from the RPE-1 cells expressing Halo-tagged DDIAS proteins in a doxycycline-inducible manner. ex., exposure. **(D)** PCR results showing homozygous knockout of the *DDIAS* gene in five independent clones in DLD-1 cells using gRNA sets A or B. Red boxes indicate the clones selected for further analysis in this study. **(E)** Western blotting of lysates from DLD-1 cells expressing Halo or Halo-tagged DDIAS proteins in a doxycycline-inducible manner. **(F)** Representative immunofluorescence images of DDIAS and γH2AX foci formation in Δ*DDIAS* DLD-1 cells expressing Halo-tagged DDIAS. Cells were treated with 50 ng/mL MMC for 24 hr where indicated. DNA was stained with DAPI. Scale bar, 10 μm.

**Figure S2.**
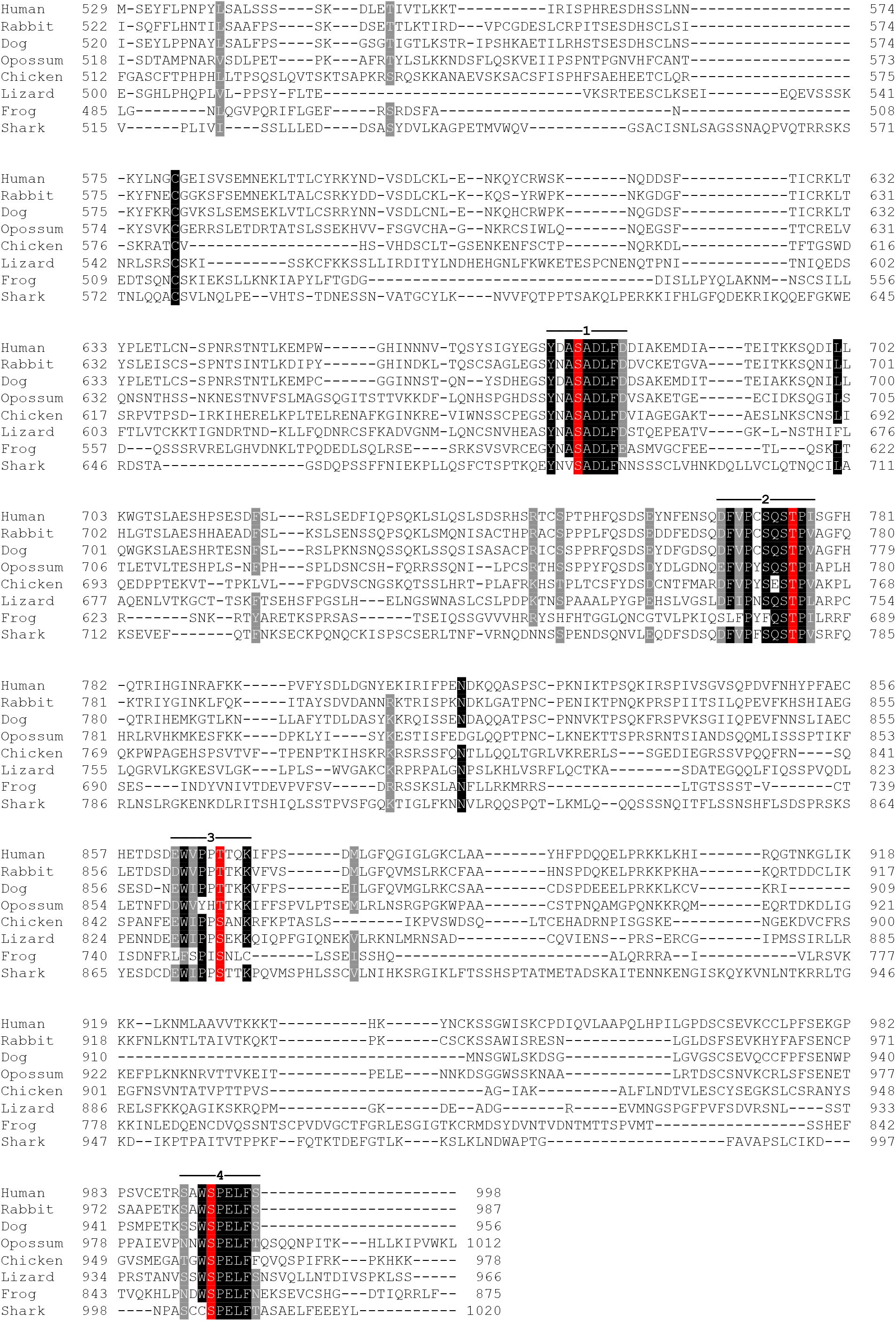
Sequence alignment of the unstructured C-terminus of vertebrate DDIAS protein orthologues, related to Figure 3. Protein sequences were aligned using Clustal Omega with conserved and similar residues highlighted in black or grey respectively using pyBoxshade. Numbered lines 1-4 denote conserved motifs synthesized as biotinylated peptides (see main text for further details). Conserved putative phosphorylation sites within these motifs are highlighted in red.

**Figure S3.**
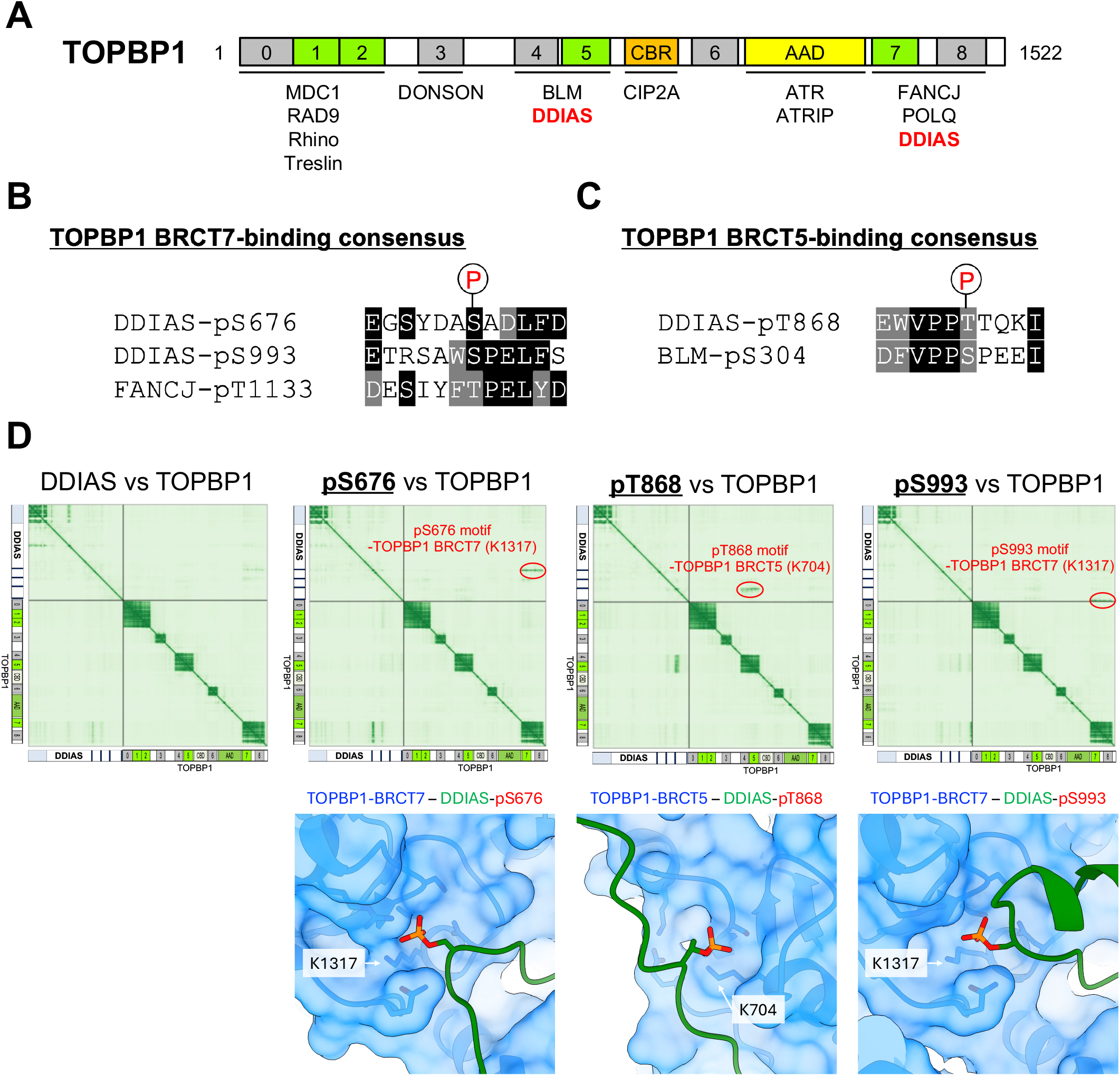
DDIAS pSer676 and pSer993 bind TOPBP1 BRCT domain 7 and DDIAS pThr868 binds TOPBP1 BRCT domain 5, related to Figure 3. **(A)** Schematic showing the layout of conserved domains and motifs in TOPBP1. Numbered boxes represent BRCT domains, with phospho-peptide binding domains in green and domains lacking phospho-peptide binding activity in grey. Names of known TOPBP1 binding partners are shown below the domains they interact with. CBR, CIP2A-binding region; AAD, ATR-activation domain. **(B)** Sequence alignments of the two TOPBP1 BRCT7-binding motifs in DDIAS and a known interaction partner, FANCJ. **(C)** Sequence alignments of the TOPBP1 BRCT5-binding motif in DDIAS and a known interaction partner, BLM. **(D)** AlphaFold 3 predictions showing conserved phospho-peptide motifs in DDIAS in complex with TOPBP1 BRCT domains.

**Figure S4.**
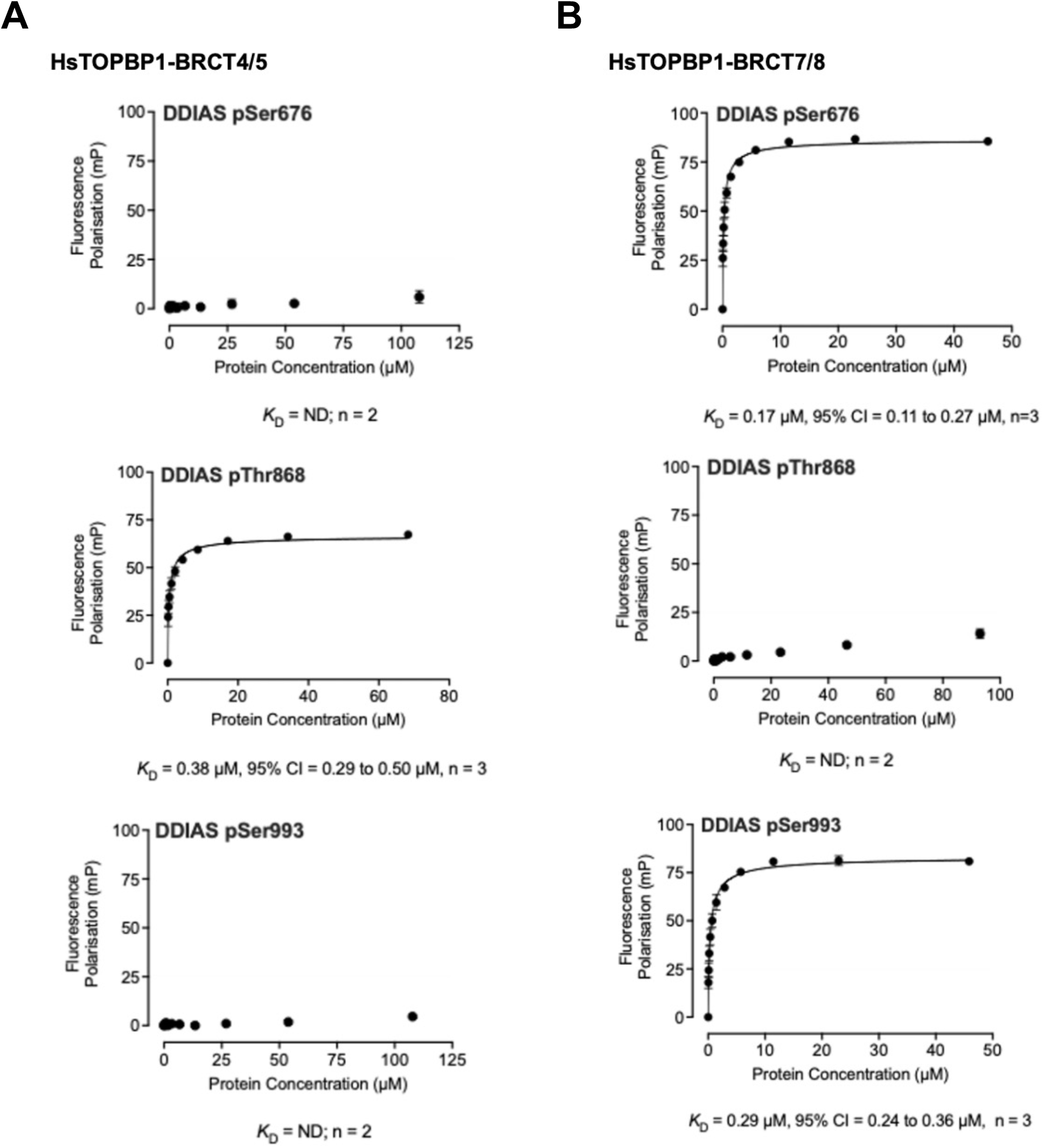
DDIAS pS676 and pS993 peptides preferentially bind to TOPBP1 tandem BRCT7+8, whereas the DDIAS pT868 peptide interacts with TOPBP1 tandem BRCT4+5, related to Figure 3. **(A)** Fluorescence polarization with recombinant TOPBP1 BRCT domains 4+5 and the indicated DDIAS phospho-peptides. **(B)** Fluorescence polarization with recombinant TOPBP1 BRCT domains 7+8 and the indicated DDIAS phospho-peptides. ND = not determined. At least two independent replicates were performed for each condition.

**Figure S5.**
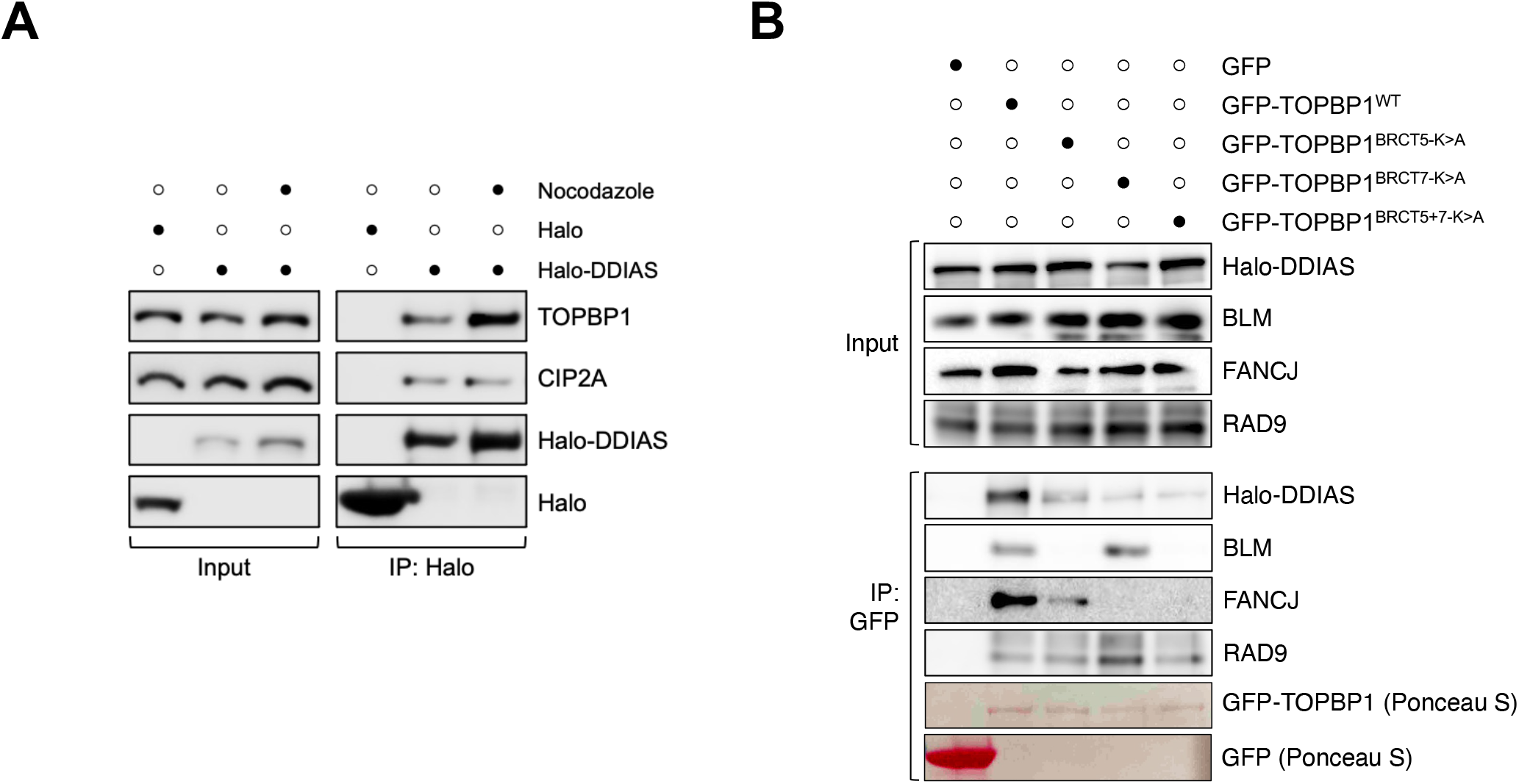
DDIAS and TOPBP1 preferentially interact in mitosis via BRCT domains 5 and 7 of TOPBP1, related to Figure 3. **(A)** Immunoprecipitations from lysates of 293FT cells transiently transfected with plasmids expressing Halo or Halo-tagged DDIAS. Where indicated, cells were treated with 100 ng/mL nocodazole for 16 hr prior to harvesting. **(B)** Immunoprecipitations from lysates of 293FT cells transiently transfected with plasmids expressing Halo-tagged DDIAS and the indicated GFP or GFP-tagged TOPBP1 proteins. Cells were treated with 100 ng/mL nocodazole for 16 h prior to harvesting.

**Figure S6.**
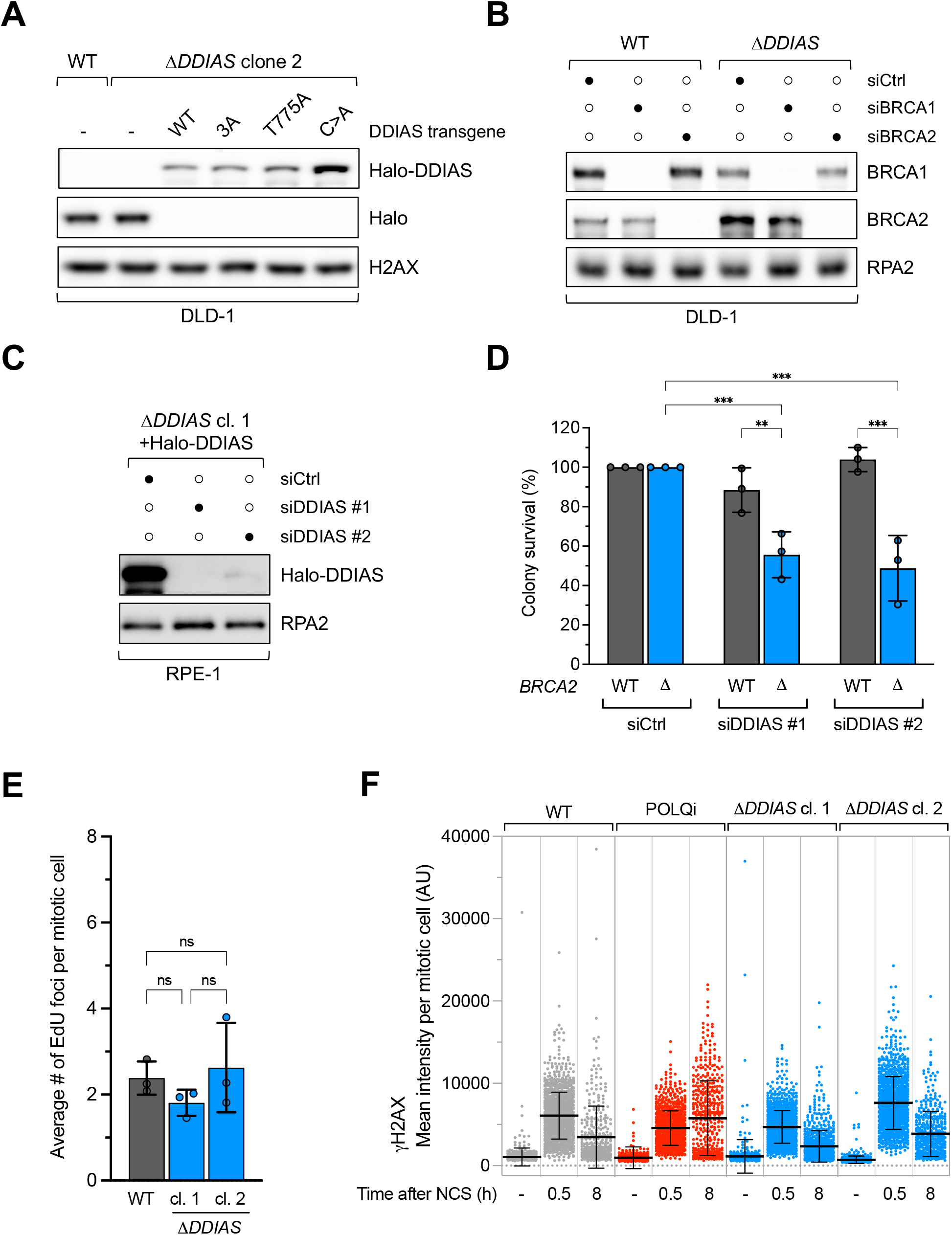
Further characterization of DDIAS-deficient cells, related to Figure 5. **(A)** Western blotting of lysates from DLD-1 cells expressing Halo or Halo-tagged DDIAS proteins in a doxycycline-inducible manner. H2AX is a loading control. **(B)** Western blotting of lysates from WT and Δ*DDIAS* DLD-1 cells treated with either control siRNA (siCtrl) or siRNAs targeting BRCA1 (siBRCA1) or BRCA2 (siBRCA2). RPA2 is a loading control. **(C)** Western blotting of lysates from Δ*DDIAS* RPE-1 cells expressing Halo-tagged DDIAS proteins and treated with either siCtrl or siRNAs targeting DDIAS (siDDIAS). **(D)** Quantification of colony survival assays with WT and Δ*BRCA2* DLD-1 cells treated with the indicated siRNAs. Error bars denote SD from n = 3 experiments. Two-way ANOVA followed by Tukey’s post hoc test was used to calculate significance. **(E)** Quantification of the number of EdU foci per mitotic cell in WT or Δ*DDIAS* DLD-1 cells treated with 400 nM APH. Each dot denotes the averages of each replicate and averages of three replicates are shown as black bars. Error bars denote SD from n = 3 experiments. One-way ANOVA followed by Tukey’s post hoc test was used to calculate significance. **(F)** Quantification of the mean intensities of γH2AX per mitotic cell, as a proxy for DSB repair in mitosis.^19^ Δ*DDIAS* DLD-1 cells and the parental DLD-1 cells were enriched for mitotic cells by pre-treatment with 9 μM CDK1 inhibitor RO-3306 for 6 hours to arrest cells in G2, followed by 100 ng/mL nocodazole treatment. Where indicated, cells were pre-treated with 10 μM POLQ inhibitor ART558; this was a positive control given that POLQ promotes DNA repair by MMEJ in mitosis, resulting in resolution of γH2AX foci.^19^ For DNA damage induction, cells were treated with 100 ng/mL NCS for 30 min and then released into fresh medium containing nocodazole for the indicated times. The mean intensities of γH2AX per mitotic cell were quantified by QIBC analyses of n = 2 replicates. Cells were gated by H3-pS10 intensity to include only mitotic cells. Averages are shown as black bars.

## REFERENCES

1. Blackford, A.N., and Jackson, S.P. (2017). ATM, ATR, and DNA-PK: The Trinity at the Heart of the DNA Damage Response. Mol. Cell 66, 801–817. 10.1016/j.molcel.2017.05.015.

2. Taylor, A.M.R., Rothblum-Oviatt, C., Ellis, N.A., Hickson, I.D., Meyer, S., Crawford, T.O., Smogorzewska, A., Pietrucha, B., Weemaes, C., and Stewart, G.S. (2019). Chromosome instability syndromes. Nature reviews. Disease primers 5, 64. 10.1038/s41572-019-0113-0.

3. Groelly, F.J., Fawkes, M., Dagg, R.A., Blackford, A.N., and Tarsounas, M. (2023). Targeting DNA damage response pathways in cancer. Nat. Rev. Cancer 23, 78–94. 10.1038/s41568-022-00535-5.

4. Blackford, A.N., and Stucki, M. (2020). How Cells Respond to DNA Breaks in Mitosis. Trends Bioch. Sci. 45, 321–331. 10.1016/j.tibs.2019.12.010.

5. Nelson, G., Buhmann, M., and Zglinicki, T.V. (2009). DNA damage foci in mitosis are devoid of 53BP1. Cell cycle (Georgetown, Tex) 8, 3379–3383. 10.4161/cc.8.20.9857.

6. Nakamura, A.J., Rao, V.A., Pommier, Y., and Bonner, W.M. (2010). The complexity of phosphorylated H2AX foci formation and DNA repair assembly at DNA double-strand breaks. Cell cycle (Georgetown, Tex) 9, 389–397. 10.4161/cc.9.2.10475.

7. Giunta, S., Belotserkovskaya, R., and Jackson, S.P. (2010). DNA damage signaling in response to double-strand breaks during mitosis. The Journal of Cell Biology 190, 197–207. 10.1083/jcb.200911156.

8. Vugt, M.A.T.M. van, Gardino, A.K., Linding, R., Ostheimer, G.J., Reinhardt, H.C., Ong, S.-E., Tan, C.S., Miao, H., Keezer, S.M., Li, J., et al. (2010). A mitotic phosphorylation feedback network connects Cdk1, Plk1, 53BP1, and Chk2 to inactivate the G(2)/M DNA damage checkpoint. PLoS biology 8, e1000287. 10.1371/journal.pbio.1000287.

9. Leimbacher, P.-A., Jones, S.E., Shorrocks, A.-M.K., Zompit, M. de M., Day, M., Blaauwendraad, J., Bundschuh, D., Bonham, S., Fischer, R., Fink, D., et al. (2019). MDC1 Interacts with TOPBP1 to Maintain Chromosomal Stability during Mitosis. Mol. Cell 74, 571–583.e8. 10.1016/j.molcel.2019.02.014.

10. Adam, S., Rossi, S.E., Moatti, N., Zompit, M.D.M., Xue, Y., Ng, T.F., Álvarez-Quilón, A., Desjardins, J., Bhaskaran, V., Martino, G., et al. (2021). The CIP2A–TOPBP1 axis safeguards chromosome stability and is a synthetic lethal target for BRCA-mutated cancer. Nat Cancer, 1–15. 10.1038/s43018-021-00266-w.

11. Laine, A., Nagelli, S.G., Farrington, C., Butt, U., Cvrljevic, A.N., Vainonen, J.P., Feringa, F.M., Grönroos, T.J., Gautam, P., Khan, S., et al. (2021). CIP2A Interacts with TopBP1 and Drives Basal-Like Breast Cancer Tumorigenesis. Cancer Res 81, 4319–4331. 10.1158/0008-5472.can-20-3651.

12. Zompit, M.D.M., Esteban, M.T., Mooser, C., Adam, S., Rossi, S.E., Jeanrenaud, A., Leimbacher, P.-A., Fink, D., Shorrocks, A.-M.K., Blackford, A.N., et al. (2022). The CIP2A-TOPBP1 complex safeguards chromosomal stability during mitosis. Nat. Commun. 13, 4143. 10.1038/s41467-022-31865-5.

13. Lin, Y.-F., Hu, Q., Mazzagatti, A., Valle-Inclán, J.E., Maurais, E.G., Dahiya, R., Guyer, A., Sanders, J.T., Engel, J.L., Nguyen, G., et al. (2023). Mitotic clustering of pulverized chromosomes from micronuclei. Nature 618, 1041–1048. 10.1038/s41586-023-05974-0.

14. Trivedi, P., Steele, C.D., Au, F.K.C., Alexandrov, L.B., and Cleveland, D.W. (2023). Mitotic tethering enables inheritance of shattered micronuclear chromosomes. Nature 618, 1049–1056. 10.1038/s41586-023-06216-z.

15. Wang, H., Qiu, Z., Liu, B., Wu, Y., Ren, J., Liu, Y., Zhao, Y., Wang, Y., Hao, S., Li, Z., et al. (2018). PLK1 targets CtIP to promote microhomology-mediated end joining. Nucleic Acids Res. 46, 10724–10739. 10.1093/nar/gky810.

16. Deng, L., Wu, R.A., Sonneville, R., Kochenova, O.V., Labib, K., Pellman, D., and Walter, J.C. (2019). Mitotic CDK Promotes Replisome Disassembly, Fork Breakage, and Complex DNA Rearrangements. Molecular Cell 73, 915–929.e6. 10.1016/j.molcel.2018.12.021.

17. Llorens-Agost, M., Ensminger, M., Le, H.P., Gawai, A., Liu, J., Cruz-García, A., Bhetawal, S., Wood, R.D., Heyer, W.-D., and Löbrich, M. (2021). POLθ-mediated end joining is restricted by RAD52 and BRCA2 until the onset of mitosis. Nat Cell Biol 23, 1095–1104. 10.1038/s41556-021-00764-0.

18. Heijink, A.M., Stok, C., Porubsky, D., Manolika, E.M., Kanter, J.K. de, Kok, Y.P., Everts, M., Boer, H.R. de, Audrey, A., Bakker, F.J., et al. (2022). Sister chromatid exchanges induced by perturbed replication can form independently of BRCA1, BRCA2 and RAD51. Nat Commun 13, 6722. 10.1038/s41467-022-34519-8.

19. Brambati, A., Sacco, O., Porcella, S., Heyza, J., Kareh, M., Schmidt, J.C., and Sfeir, A. (2023). RHINO directs MMEJ to repair DNA breaks in mitosis. Science 381, 653–660. 10.1126/science.adh3694.

20. Gelot, C., Kovacs, M.T., Miron, S., Mylne, E., Haan, A., Boeffard-Dosierre, L., Ghouil, R., Popova, T., Dingli, F., Loew, D., et al. (2023). Polθ is phosphorylated by PLK1 to repair double-strand breaks in mitosis. Nature 621, 415–422. 10.1038/s41586-023-06506-6.

21. Tsukada, K., Jones, S.E., Bannister, J., Durin, M.-A., Vendrell, I., Fawkes, M., Fischer, R., Kessler, B.M., Chapman, J.R., and Blackford, A.N. (2024). BLM and BRCA1-BARD1 coordinate complementary mechanisms of joint DNA molecule resolution. Mol. Cell 84, 640–658.e10. 10.1016/j.molcel.2023.12.040.

22. Germann, S.M., Schramke, V., Pedersen, R.T., Gallina, I., Eckert-Boulet, N., Oestergaard, V.H., and Lisby, M. (2014). TopBP1/Dpb11 binds DNA anaphase bridges to prevent genome instability. The Journal of Cell Biology 204, 45–59. 10.1083/jcb.201305157.

23. Blackford, A.N., Nieminuszczy, J., Schwab, R.A., Galanty, Y., Jackson, S.P., and Niedzwiedz, W. (2015). TopBP1 Interacts with BLM to Maintain Genome Stability but Is Dispensable for Preventing BLM Degradation. Mol. Cell 57, 1133–1141. 10.1016/j.molcel.2015.02.012.

24. Broderick, R., Nieminuszczy, J., Blackford, A.N., Winczura, A., and Niedzwiedz, W. (2015). TOPBP1 recruits TOP2A to ultra-fine anaphase bridges to aid in their resolution. Nat. Commun. 6, 6572. 10.1038/ncomms7572.

25. Pedersen, R.T., Kruse, T., Nilsson, J., Oestergaard, V.H., and Lisby, M. (2015). TopBP1 is required at mitosis to reduce transmission of DNA damage to G1 daughter cells. The Journal of Cell Biology 210, 565–582. 10.1083/jcb.201502107.

26. Fleury, H., MacEachern, M.K., Stiefel, C.M., Anand, R., Sempeck, C., Nebenfuehr, B., Maurer-Alcalá, K., Ball, K., Proctor, B., Belan, O., et al. (2023). The APE2 nuclease is essential for DNA double-strand break repair by microhomology-mediated end joining. Mol. Cell 83, 1429–1445.e8. 10.1016/j.molcel.2023.03.017.

27. Bagge, J., Petersen, K.V., Karakus, S.N., Nielsen, T.M., Rask, J., Brøgger, C.R., Jensen, J., Skouteri, M., Carr, A.M., Hendriks, I.A., et al. (2025). TopBP1 coordinates DNA repair synthesis in mitosis via recruitment of the nuclease scaffold SLX4. Commun. Biol. 8, 1005. 10.1038/s42003-025-08442-9.

28. Martin, P.R., Nieminuszczy, J., Kozik, Z., Jakub, N., Kowalski, S., Lecot, M., Vorhauser, J., Lane, K.A., Kanellou, A., Mansfeld, J., et al. (2025). The CIP2A-TOPBP1 axis facilitates mitotic DNA repair via MiDAS and MMEJ. Nat. Commun. 16, 10623. 10.1038/s41467-025-65594-2.

29. Haan, L. de, Dijt, S.J., García-López, A., Ruan, D., Martzios, P., Bakker, F.J., Everts, M., Warner, H., Mol, F.N., Chapman, J.R., et al. (2025). CIP2A mediates mitotic recruitment of SLX4/MUS81/XPF to resolve replication stress-induced DNA lesions. Nat. Commun. 17, 13. 10.1038/s41467-025-66549-3.

30. Meroni, A., Pellizzari, A., Varisco, N., Leone, F., Greco, G., Hänel, A., Brasier-Lutz, P., Witzel, I., Sartori, A.A., and Stucki, M. (2026). CIP2A recruits SLX4-MUS81-XPF in mitosis and protects against replication stress. EMBO Rep., 1–19. 10.1038/s44319-026-00807-3.

31. Nakaya, N., Hemish, J., Krasnov, P., Kim, S.-Y., Stasiv, Y., Michurina, T., Herman, D., Davidoff, M.S., Middendorff, R., and Enikolopov, G. (2007). noxin, a Novel Stress-Induced Gene Involved in Cell Cycle and Apoptosis. Mol. Cell. Biol. 27, 5430–5444. 10.1128/mcb.00551-06.

32. Brunette, G.J., Jamalruddin, M.A., Baldock, R.A., Clark, N.L., and Bernstein, K.A. (2019). Evolution-based screening enables genome-wide prioritization and discovery of DNA repair genes. Proceedings of the National Academy of Sciences 116, 19593–19599. 10.1073/pnas.1906559116.

33. Rogakou, E.P., Boon, C., Redon, C., and Bonner, W.M. (1999). Megabase chromatin domains involved in DNA double-strand breaks in vivo. The Journal of Cell Biology 146, 905–916.

34. Abramson, J., Adler, J., Dunger, J., Evans, R., Green, T., Pritzel, A., Ronneberger, O., Willmore, L., Ballard, A.J., Bambrick, J., et al. (2024). Accurate structure prediction of biomolecular interactions with AlphaFold 3. Nature 630, 493–500. 10.1038/s41586-024-07487-w.

35. Day, M., Oliver, A.W., and Pearl, L.H. (2021). Phosphorylation-dependent assembly of DNA damage response systems and the central roles of TOPBP1. DNA Repair 108, 103232. 10.1016/j.dnarep.2021.103232.

36. Leung, C.C.Y., Gong, Z., Chen, J., and Glover, J.N.M. (2011). Molecular Basis of BACH1/FANCJ Recognition by TopBP1 in DNA Replication Checkpoint Control. The Journal of biological chemistry 286, 4292–4301. 10.1074/jbc.m110.189555.

37. Sun, L., Huang, Y., Edwards, R.A., Yang, S., Blackford, A.N., Niedzwiedz, W., and Glover, J.N.M. (2017). Structural Insight into BLM Recognition by TopBP1. Structure 25, 1582–1588.e3. 10.1016/j.str.2017.08.005.

38. Elia, A.E.H., Cantley, L.C., and Yaffe, M.B. (2003). Proteomic Screen Finds pSer/pThr-Binding Domain Localizing Plk1 to Mitotic Substrates. Science 299, 1228–1231. 10.1126/science.1079079.

39. Lens, S.M.A., Voest, E.E., and Medema, R.H. (2010). Shared and separate functions of polo-like kinases and aurora kinases in cancer. Nat. Rev. Cancer 10, 825–841. 10.1038/nrc2964.

40. Vassilev, L.T., Tovar, C., Chen, S., Knezevic, D., Zhao, X., Sun, H., Heimbrook, D.C., and Chen, L. (2006). Selective small-molecule inhibitor reveals critical mitotic functions of human CDK1. Proc. Natl. Acad. Sci. 103, 10660–10665. 10.1073/pnas.0600447103.

41. Steegmaier, M., Hoffmann, M., Baum, A., Lénárt, P., Petronczki, M., Krssák, M., Gürtler, U., Garin-Chesa, P., Lieb, S., Quant, J., et al. (2007). BI 2536, a potent and selective inhibitor of polo-like kinase 1, inhibits tumor growth in vivo. Current Biology 17, 316–322. 10.1016/j.cub.2006.12.037.

42. Schou, K.B., Mandacaru, S., Tahir, M., Tom, N., Nilsson, A.-S., Andersen, J.S., Tiberti, M., Papaleo, E., and Bartek, J. (2024). Exploring the structural landscape of DNA maintenance proteins. Nat. Commun. 15, 7748. 10.1038/s41467-024-49983-7.

43. Cong, K., and Cantor, S.B. (2022). Exploiting replication gaps for cancer therapy. Mol. Cell 82, 2363–2369. 10.1016/j.molcel.2022.04.023.

44. Ramirez-Otero, M.A., and Costanzo, V. (2024). “Bridging the DNA divide”: Understanding the interplay between replication-gaps and homologous recombination proteins RAD51 and BRCA1/2. DNA Repair 141, 103738. 10.1016/j.dnarep.2024.103738.

45. Liu, L., Michowski, W., Kolodziejczyk, A., and Sicinski, P. (2019). The cell cycle in stem cell proliferation, pluripotency and differentiation. Nat. Cell Biol. 21, 1060–1067. 10.1038/s41556-019-0384-4.

46. Zimmer, J., Tacconi, E.M.C., Folio, C., Badie, S., Porru, M., Klare, K., Tumiati, M., Markkanen, E., Halder, S., Ryan, A., et al. (2016). Targeting BRCA1 and BRCA2 Deficiencies with G-Quadruplex-Interacting Compounds. Molecular Cell 61, 449–460. 10.1016/j.molcel.2015.12.004.

47. Schindelin, J., Arganda-Carreras, I., Frise, E., Kaynig, V., Longair, M., Pietzsch, T., Preibisch, S., Rueden, C., Saalfeld, S., Schmid, B., et al. (2012). Fiji: an open-source platform for biological-image analysis. Nature Methods 9, 676–682. 10.1038/nmeth.2019.

48. Cescutti, R., Negrini, S., Kohzaki, M., and Halazonetis, T.D. (2010). TopBP1 functions with 53BP1 in the G1 DNA damage checkpoint. The EMBO Journal 29, 3723–3732. 10.1038/emboj.2010.238.

49. Shorrocks, A.-M.K., Jones, S.E., Tsukada, K., Moron, C.A., Belblidia, Z., Shen, J., Vendrell, I., Fischer, R., Kessler, B.M., and Blackford, A.N. (2021). The Bloom syndrome complex senses RPA-coated single-stranded DNA to restart stalled replication forks. Nat. Commun. 12, 585. 10.1038/s41467-020-20818-5.

50. Becker, J.R., Cuella-Martin, R., Barazas, M., Liu, R., Oliveira, C., Oliver, A.W., Bilham, K., Holt, A.B., Blackford, A.N., Heierhorst, J., et al. (2018). The ASCIZ-DYNLL1 axis promotes 53BP1-dependent non-homologous end joining and PARP inhibitor sensitivity. Nat. Commun. 9, 5406. 10.1038/s41467-018-07855-x.

51. Ran, F.A., Hsu, P.D., Wright, J., Agarwala, V., Scott, D.A., and Zhang, F. (2013). Genome engineering using the CRISPR-Cas9 system. Nature protocols 8, 2281–2308. 10.1038/nprot.2013.143.

52. Hertzog, J., Junior, A.G.D., Rigby, R.E., Donald, C.L., Mayer, A., Sezgin, E., Song, C., Jin, B., Hublitz, P., Eggeling, C., et al. (2018). Infection with a Brazilian isolate of Zika virus generates RIG-I stimulatory RNA and the viral NS5 protein blocks type I IFN induction and signaling. Eur. J. Immunol. 48, 1120–1136. 10.1002/eji.201847483.

53. Zatreanu, D., Robinson, H.M.R., Alkhatib, O., Boursier, M., Finch, H., Geo, L., Grande, D., Grinkevich, V., Heald, R.A., Langdon, S., et al. (2021). Polθ inhibitors elicit BRCA-gene synthetic lethality and target PARP inhibitor resistance. Nat Commun 12, 3636. 10.1038/s41467-021-23463-8.

54. Chen, X., Ali, Y.I., Fisher, C.E., Arribas-Bosacoma, R., Rajasekaran, M.B., Williams, G., Walker, S., Booth, J.R., Hudson, J.J., Roe, S.M., et al. (2021). Uncovering an allosteric mode of action for a selective inhibitor of human Bloom syndrome protein. Elife 10, e65339. 10.7554/elife.65339.

55. Hannus, M., Beitzinger, M., Engelmann, J.C., Weickert, M.-T., Spang, R., Hannus, S., and Meister, G. (2014). siPools: highly complex but accurately defined siRNA pools eliminate off-target effects. Nucleic Acids Research. 10.1093/nar/gku480.

